# A biphasic metabolic–growth trade-off governs β-lactam inoculum effects

**DOI:** 10.1101/2025.10.14.682300

**Authors:** Ariane R. Kalifa Torjeman, Kielan Singh, Philopatier Ibrahim, Trent Moulder, Tiffany Rakela, Yabesera Negussie, Hansika Vusikamalla, Estefania Marin Meneses, Maximiliano Barbosa Mendoza, Debora Pereira Mcmenamin, Daniel Winget, Taija White, Archit Pipalva, Naziba Nuha, Julia Gumirov, Allison. J. Lopatkin, Robert P. Smith

## Abstract

The inoculum effect (IE) reduces antibiotic efficacy *in vitro* and in clinical settings, yet remains difficult to predict. IE limits antibiotic efficacy but remains unpredictable, particularly in resistant bacteria, where resistance mechanisms and bacterial physiology interact in poorly understood ways. Using *Escherichia coli* expressing the NDM-1 β-lactamase and clinical isolates, we quantified metabolism, growth rate, β-lactamase expression, and IE across diverse growth environments. Across enzyme classes, antibiotics, and clinical isolates, we find that IE is governed by a conserved, biphasic dependence on metabolism normalized by growth rate. Mathematical modeling shows that this behavior reflects a trade⍰off between β⍰lactamase⍰mediated protection and metabolism⍰potentiated antibiotic lethality. These findings establish a predictive, physiology-based framework for IE in resistant bacteria and explain why resistance determinants alone fail to predict treatment outcomes across environments, including the clinic.

**Significance Statement:** Antibiotic efficacy depends on both bacterial resistance and physiology, yet these factors are rarely integrated when predicting treatment outcomes. One important consequence of their interaction is the inoculum effect (IE), in which antibiotic efficacy depends on the density of a resistant bacterial population. Here, we show that IE in β⍰lactamase–expressing bacteria is governed by a conserved physiological trade⍰off between metabolism and growth. Across growth environments, antibiotics, inocula, and clinical isolates, IE is strongest at intermediate metabolic states, reflecting a balance between resistance⍰mediated protection and metabolism⍰potentiated antibiotic lethality. This framework helps explain why resistance determinants alone are insufficient to account for IE and underscores the role of bacterial physiology in shaping antibiotic responses.

## Introduction

It has long been recognized that bacterial population density can strongly reduce antibiotic efficacy. This phenomenon, called the inoculum effect (IE), has been observed for nearly all bacteria and antibiotics (1-4). At a given antibiotic concentration, a high-density population will survive antibiotic treatment and grow. However, if the population density is sufficiently low, the population is susceptible to the antibiotic and dies. Reports from the clinic have demonstrated that IE can increase morbidity and mortality by reducing antibiotic efficacy (5-7) and can promote the evolution of additional resistance mechanisms (8, 9). Despite being first reported in 1949 (10), no reliable framework exists to predict or mitigate IE in clinical settings.

Antibiotic efficacy is increasingly understood to depend on bacterial physiological state, including metabolism and growth rate. Multiple studies have shown that increased bacterial metabolism increases antibiotic lethality (11), while bacterial growth has been shown to increase (12) or decrease (13) antibiotic lethality. Recent work demonstrated that, across multiple Gram⍰negative bacteria and antibiotics, IE emerges from interactions between metabolism and growth rate (14, 15). In bacteria lacking enzyme-mediated resistance, increasing bacterial metabolism relative to growth rate decreases IE (16). Mechanistically, populations initiated at high density experience only a brief period of logarithmic growth, when metabolism and antibiotic lethality are maximal, before entering the stationary phase. In contrast, low⍰density populations remain metabolically active for longer, rendering them more susceptible to metabolism⍰potentiated antibiotic killing. Differences in the time each population spends in this high metabolic state determine the magnitude of IE (16).

This physiological framework appears to be in tension with a large body of work implicating β-lactamase expression as a major determinant of IE for β-lactam antibiotics (3, 5). β-lactamases inactivate β-lactam antibiotics, conferring resistance (17), and IE in β-lactamase-expressing bacteria has been associated with antibiotic failure and increased patient mortality (18-20). Interestingly, expression of heterologous proteins, including β-lactamases, imposes metabolic costs that can alter [ATP] (21, 22) and reduce growth rate (23, 24). Thus, β-lactamase expression can both protect cells from antibiotics and reshape the metabolic and growth dynamics previously shown to govern IE.

One previously proposed explanation for IE in β-lactamase-expressing bacteria is collective antibiotic inactivation, where higher-density populations degrade greater amounts of antibiotic (Fig. 1A)(25-29). While this mechanism may explain IE under some conditions, it does not broadly predict IE. Many β-lactamase-positive strains fail to exhibit IE (30, 31), and IE strength varies widely across β-lactam antibiotics and clinical isolates (32). Clinical studies further demonstrate that β-lactamase-positive isolates do not consistently exhibit IE, even when controlling for bacterial species, enzyme type, and antibiotic (5). Together, these observations indicate that resistance determinants alone are insufficient to predict IE and suggest that higher-order physiological constraints shape inoculum-dependent responses to β-lactams.

**Figure 1:**
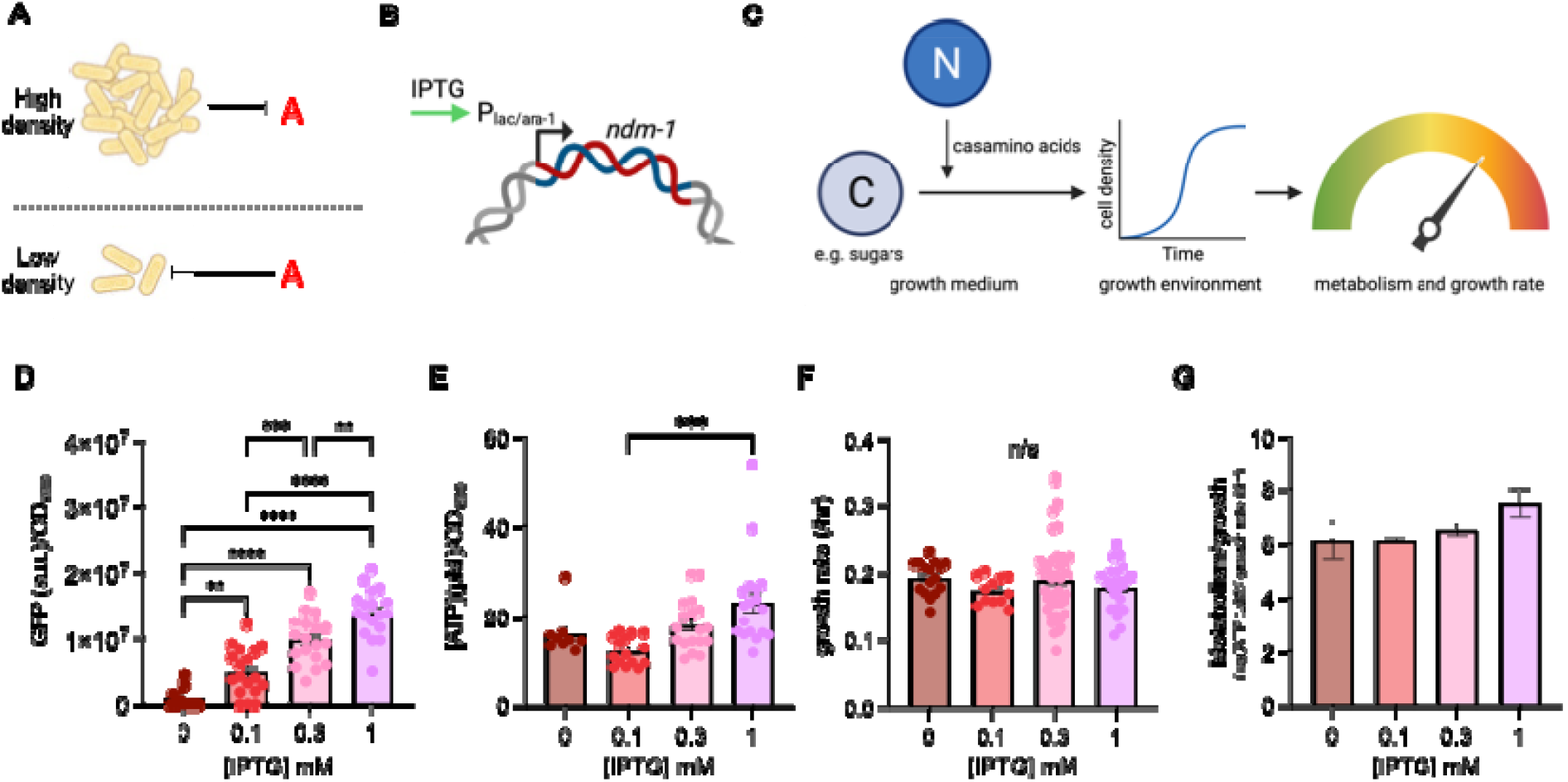
Orthogonal manipulation of β□lactamase expression, metabolism, and growth in a reductionist *E. coli* system. A) Schematic illustration of the inoculum effect (IE). At high initial density, bacterial populations resist antibiotic treatment and grow (top), whereas populations initiated at low density are susceptible and die (bottom). B) *E. coli* expressing the NDM⍰1 β⍰lactamase from a plasmid under control of an IPTG-inducible P_lac/ara-1_ promoter. C) Experimental framework used to manipulate bacterial metabolism ([ATP]) and growth rate. The carbon source defined the growth environment, while varying the nitrogen source (casamino acids) modulated metabolism and growth within a given environment. D) Effect of increasing IPTG on downstream expression. GFP (arbitrary units, a.u.) was normalized by cell density (OD_600_) and measured after 5□h of induction, confirming inducible and dose⍰dependent expression. ANOVA, P < 0.0001, Tukey HSD (** P ≤ 0.0031, *** P = 0.0002, **** P < 0.0001). For panels D-G, the standard error of the mean (SEM) is calculated from ≥ 3 biological replicates per % casamino acids. Similar trends in GFP induction were observed at 3, 7, and 9 hours post-induction (Supplementary Fig. 1). E) Effect of increasing [IPTG] on cellular [ATP], normalized by OD_600_. ANOVA, P = 0.0004, Tukey HSD, *** P = 0.0002. F) Growth rate as a function of [IPTG]. No significant differences were observed across IPTG conditions (n/s). Growth curves are in Supplementary Fig. 1; average residuals for curve fitting in Supplementary Table 1. G) Metabolism normalized by growth as a function of increasing [IPTG]. [ATP] and growth rate from panels E and F.

Here, we test whether interactions among key physiological predictors of antibiotic efficacy, metabolism and growth rate, and β-lactamase expression give rise to general and predictable patterns of IE strength beyond those driven solely by β-lactam resistance. Addressing this question may help define growth environments in which IE is amplified or suppressed, reconcile conflicting reports on the clinical importance of IE (33), and provide a physiology⍰based framework for interpreting antibiotic efficacy.

## Results

### A reductionist experimental framework to study β⍰ lactamase⍰mediated inoculum effects

To understand how bacterial metabolism, growth rate, and β-lactamase expression would determine IE, we used a strain of *Escherichia coli* that produces the NDM-1 β-lactamase (34) from a plasmid under the regulation of an isopropyl-β-D-thiogalactopyranoside (IPTG) inducible promoter, P_lac/ara-1_(35) (Fig. 1B). We chose to use the NDM-1 β-lactamase given its clinical relevance, global distribution, and ability to inactivate a wide range of β-lactams (36). Introducing an inducible copy of NDM-1 into a laboratory strain of *E. coli* allowed us to determine how β-lactamase expression reshapes metabolism, growth, and IE without confounding effects from strain-specific mutations, evolutionary history, or additional resistance mechanisms present in clinical isolates.

Previous work has shown that the type of carbon source can influence both metabolism (37) and growth rate (38). Accordingly, to systematically vary metabolism and growth, we grew bacteria in a minimal medium containing a carbon source, which defined the growth environment. We also provided casamino acids as a nitrogen source (Fig. 1C). For each carbon source, we supplied three casamino acid concentrations (0.1, 0.5, and 1%). By averaging across nitrogen concentrations, we identified a robust, carbon-source-dependent effect on bacterial metabolism and growth rate, yielding a single value for each growth environment. To vary NDM-1 orthogonally to the growth environment, we provided different [IPTG] (0, 0.1, 0.3, and 1 mM) in the growth medium, an approach previously shown to alter the expression of elements downstream of the P_lac-ara-1_ promoter (35).

While multiple facets of bacterial metabolism can be measured, we focused on ATP as an integrative proxy of metabolic state, as it correlates with other metabolic measures such as the NAD^+^/NADH ratio and oxygen consumption(14, 16), without assigning causality to specific pathways. To measure [ATP], we used a previously published method that quantifies bacterial metabolism independent of changes in growth rate (14). To measure the growth rate, we obtained high-resolution cell density (OD_600_) measurements in a microplate reader without antibiotics. We then used a custom-built model to estimate the maximum growth rate by fitting a logistic growth curve through the resulting experimental growth curve(16). Finally, to measure β-lactamase expression, we replaced NDM-1 with a green fluorescent protein (GFP); GFP was driven by the same promoter using the same plasmid backbone, serving as a reliable surrogate for NDM-1 expression. Together, this approach allowed us to measure bacterial metabolism (ATP), growth rate, and β-lactamase expression for a given growth environment.

To start, we measured the effect of increasing [IPTG] on GFP expression. When grown in M9 medium with glucose as a carbon source, we found that as [IPTG] increased, GFP per cell increased significantly, confirming the inducible and dose-dependent nature of our expression system (Fig. 1D). Next, we measured [ATP] normalized by cell density in medium with increasing [IPTG] in our NDM-1-expressing strain and found that at 1□mM IPTG, [ATP] per cell was significantly greater compared to the 0.1□mM IPTG condition (Fig. 1E). In contrast, we observed no significant differences in growth rate across [IPTG] conditions (Fig. 1F). Thus, increasing [IPTG] selectively altered metabolic output without affecting growth. Finally, we normalized log⍰transformed [ATP] by growth rate for each [IPTG] to account for differences in scale between [ATP] and growth rate; a metric hereafter referred to as metabolism normalized by growth, a composite physiological predictor. Metabolism normalized by growth reflects the balance between metabolism⍰potentiated antibiotic lethality and growth⍰rate–dependent exit from log⍰phase growth, which together mechanistically govern the IE (16). We found that increasing [IPTG] generally led to higher metabolism normalized by growth (Fig. 1G). Because ATP and growth rate were measured in independent experimental sets, metabolism normalized by growth values are presented as descriptive metrics derived from pooled means rather than as quantities subjected to direct statistical testing (see Methods). Taken together, [IPTG] increased GFP expression while nonlinearly modulating metabolism and growth, providing an orthogonal platform to test how metabolism, growth, and β⍰lactamase expression shape the inoculum effect.

### Model⍰predicted effects of β ⍰lactamase expression, metabolism, and growth on the inoculum effect

To guide our experiments and provide insight into how growth and metabolism impact IE, we developed a simple mathematical model (see *Methods*, Eq.□1). The model balances logistic growth against antibiotic-induced death, which is reduced by β-lactamase expression in a density-dependent manner. Under this framework, IE arises from the balance between growth and metabolism, while selectively increasing β-lactamase expression protects high-density populations. We first simulated populations initiated at high and low densities while increasing β-lactamase expression, holding metabolism and growth constant. The model predicted that increasing β-lactamase expression increases IE, quantified as ΔMIC, the difference in the minimum inhibitory concentration (MIC) between high- and low-density populations (Fig.□2A and B). Increasing β-lactamase expression expanded the antibiotic range over which only high-density populations survived, thereby increasing ΔMIC. We experimentally tested this prediction by measuring ΔMIC in NDM-1-expressing *E*.□*coli* grown with either 0 or 0.1□mM IPTG, conditions that increased β-lactamase expression without altering [ATP] or growth rate (Fig.□2C and D). To determine ΔMIC, we challenged NDM-1-expressing bacteria with the β-lactam antibiotic ampicillin, as it is often used to develop mechanisms of antibiotic resistance (39). We used two initial bacterial densities, 10^5^ (5.3 x 10^5^ ± 1.24 x 10^4^ CFU/mL) and 10^4^ (2.34 x 10^4^ ± 3.16 x 10^3^ □CFU/mL), both of which showed similar GFP expression rates, indicating that initial cell density did not affect expression from the P_lac-ara-1_ promoter (Supplementary Fig.□2). Consistent with model predictions, ΔMIC was significantly greater in the induced condition (Fig.□2D), demonstrating that increased β - lactamase expression increases IE absent changes in metabolism and growth. Next, simulations holding β-lactamase expression constant predicted that increasing metabolism relative to growth reduces ΔMIC (Fig.□2E and F). Experimentally, growth with adenine as the carbon source increased metabolism, normalized by growth, without altering β-lactamase expression. Together, this resulted in a reduced ΔMIC compared with glucose (Fig. 2G and H), demonstrating that increasing metabolism, normalized by growth, in the absence of changes in β-lactamase expression, decreases ΔMIC. Thus, the model accurately predicts both β-lactamase- and metabolism-driven modulation of IE. Importantly, we found that while GFP expression did not generally affect [ATP] or growth rate compared with NDM-1 expression, only NDM-1 expression increased ΔMIC as a function of [IPTG] (Supplementary Fig.□3). Although GFP expression could alter [ATP] and growth, it did not confer ampicillin resistance and did not give rise to changes in IE. Together, these results demonstrate that changes in ΔMIC and IE were specific to β-lactamase activity, rather than a generic consequence of heterologous protein expression. Importantly, this framework predicts that resistance mechanisms shape IE only within physiological bounds set by metabolism and growth.

**Figure 2:**
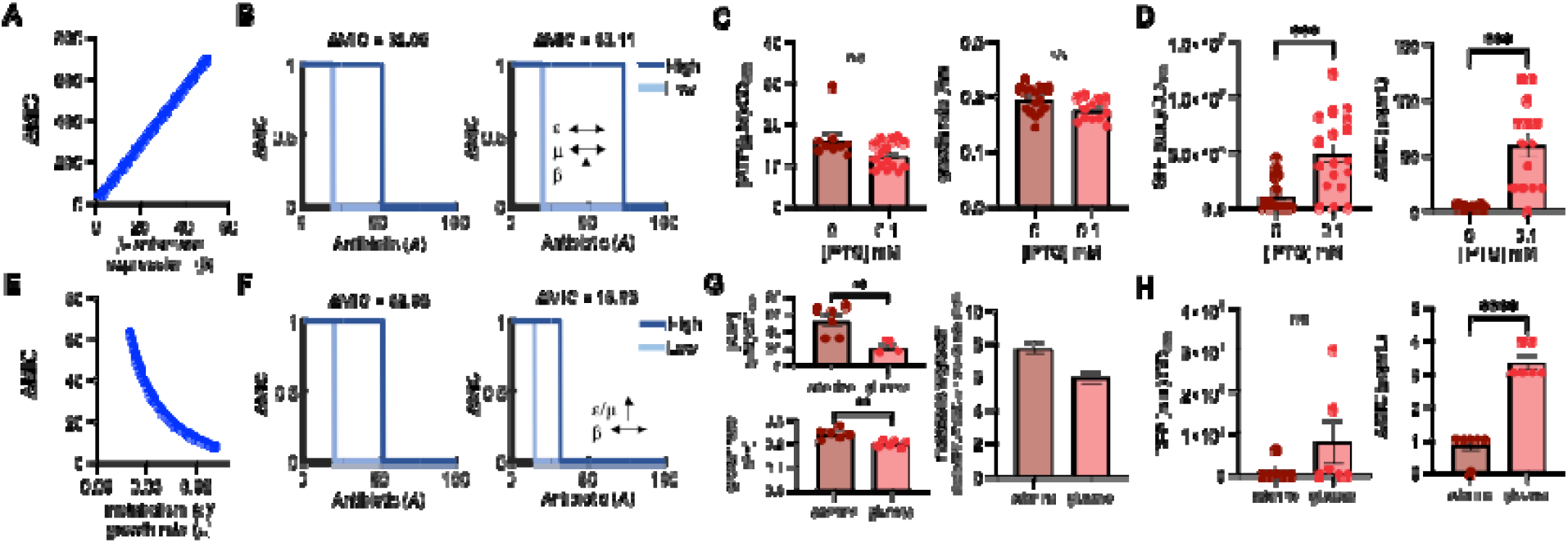
A simple mathematical model predicts experimentally verified changes to ΔMIC. A) Simulations predict that increasing β-lactamase expression (*β*) increases ΔMIC. For all simulations, *t* = 24 hours. Justification of parameters in *Methods*; values used for simulations in Supplementary Table 2. B) Increasing β-lactamase (*β*) expression from 1 (left) to 3 (right) increases ΔMIC. Metabolism (*ε*) and growth (*μ*) are kept constant. C) Experiment: Left: [ATP] normalized by cell density for *E. coli* grown in medium with glucose and with either 0 or 0.1 mM IPTG. Right: Growth rate for *E. coli* grown in medium with glucose and with either 0 or 0.1 mM IPTG. For both panels, SEM from ≥ 3 biological replicates. n/s = not significant. D) Experiments: Left: GFP normalized by cell density for *E. coli* grown in medium with glucose and with either 0 or 0.1 mM IPTG. *** Welch’s t-test, P = 0.0002. Right: ΔMIC *E. coli* grown in medium with glucose and with either 0 or 0.1 mM IPTG. *** Welch’s t-test, P = 0.0003. For both panels, SEM from 6 biological replicates. Raw MIC data in Supplementary Fig. 3. E) Simulations predict that increasing metabolism (*ε*) / growth rate (μ) decreases ΔMIC. F) Increasing metabolism (*ε*) / growth rate (μ) from 0.033 (left) to 0.05 (right) decreases ΔMIC. G) Experiment: Left, top: [ATP] normalized by cell density for *E. coli* grown in medium with adenine or glucose and 1% casamino acids (CAA) with 0 mM IPTG. ** Welch’s t-test, P = 0.0072. Left, bottom: Growth rate for *E. coli* grown in medium with adenine or glucose and 1% CAA with 0 mM IPTG. ** Welch’s t-test, P = 0.0042. For both panels, SEM from ≥ 3 biological replicates. Right: [ATP]/growth rate determined from left panels. H) Experiments: Left: GFP normalized by cell density for *E. coli* grown in medium with adenine or glucose and with 0 mM IPTG. Right: ΔMIC *E. coli* grown in medium with adenine or glucose and with 0 IPTG. For both panels, SEM from 6 biological replicates. **** Welch’s t-test, P < 0.0001.

### Model and experiments reveal a biphasic dependence of ΔMIC on metabolism and growth

Using our model, we determined ΔMIC across a range of metabolism and β-lactamase expression values while keeping the growth rate constant (Fig. 3A). We found that combinations of β-lactamase expression and metabolism yielded a broad range of ΔMIC values. Slices through the simulated landscape provided insight into the relationship between metabolism and β-lactamase expression on ΔMIC (Fig. 3B). Our model predicts that if β-lactamase expression increases relatively faster than metabolism, ΔMIC exhibits a biphasic dependence on metabolism normalized by growth rate. Specifically, ΔMIC initially increases with increasing metabolism/growth rate, but at higher values, further increases in metabolism/growth rate reduce MIC, reflecting increased metabolism-and potentiated antibiotic lethality. The relative scaling of β-lactamase expression governs the prominence of this peak, such that larger increases in β-lactamase expression relative to metabolism produce a more pronounced biphasic response. In contrast, if metabolism increases faster than β-lactamase expression, MIC decreases monotonically with increasing metabolism/growth rate. These trends were consistent across a wide range of model parameters (Supplementary Fig. 4), demonstrating the robustness of these predictions. Importantly, β-lactamase expression alone is insufficient to account for the biphasic relationship, which emerges only in conjunction with changes in metabolism and growth rate (Supplementary Fig. 4).

**Figure 3:**
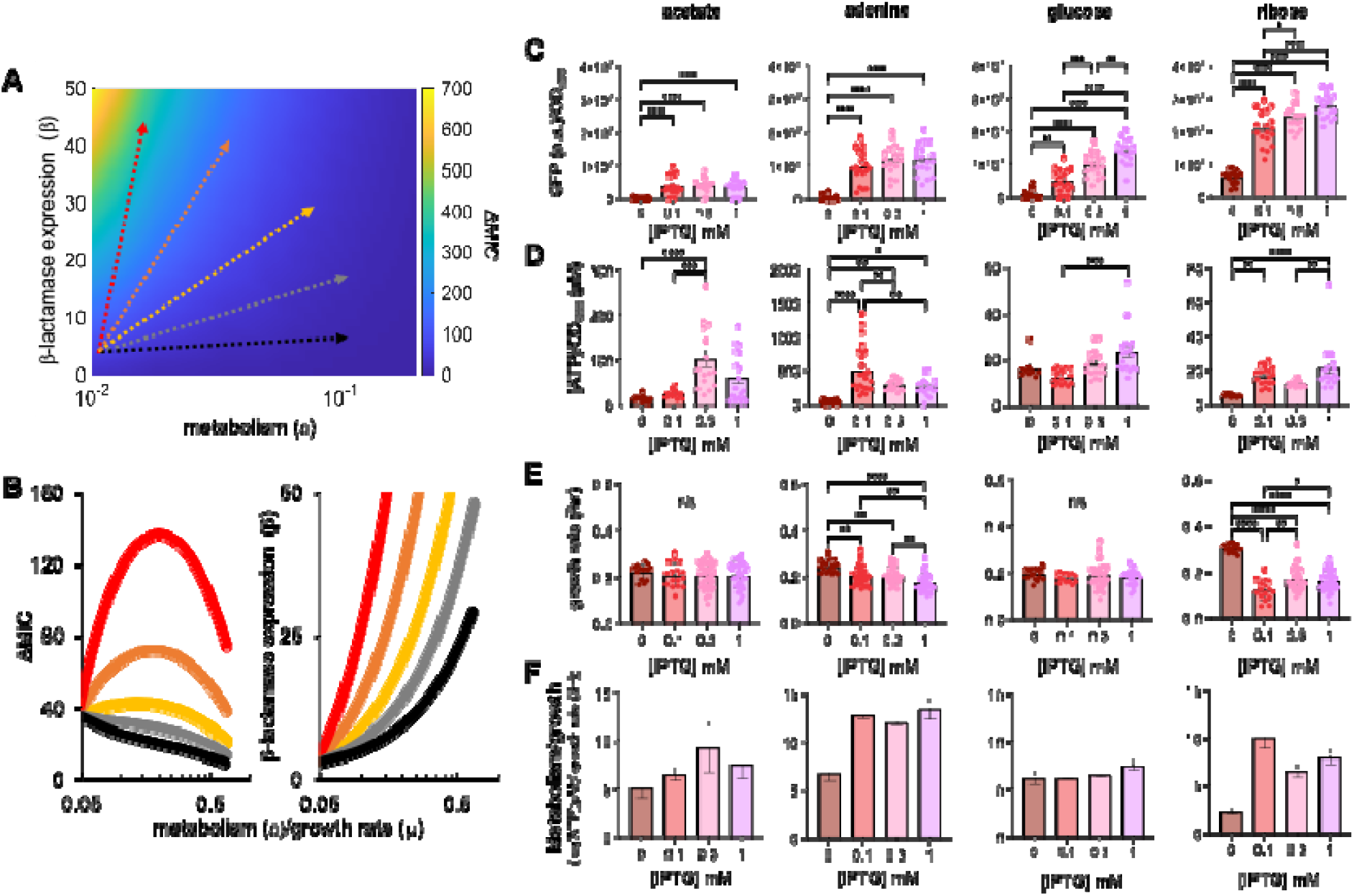
A metabolism-growth landscape reveals a biphasic relationship between ΔMIC, metabolism, and growth rate. **A)** Simulated heat map showing the effect of increasing metabolism (*ε*) and β-lactamase expression (*β*) on ΔMIC. Growth rate (*μ*) held constant. The arrows indicate an approximate slice of the heat map, revealing relationships between ε/μ and ΔMIC for different values of β, as shown in panel B. Sensitivity analysis, including the removal of ε from the *per capita* death rate term (Eq. 1), is presented in Supplementary Fig. 4. **B)** Left: Simulations showing the effect of increasing metabolism (*ε*) and β-lactamase expression (β) on the relationship with ΔMIC. Colors match arrows in panel A. Right: Simulations showing the value of β-lactamase expression (β) for each value of metabolism (*ε*)/growth rate (*μ*) in the left panel. **C)** GFP expression in mid-log (5 hours) in *E. coli* grown in M9 medium with different carbon sources and [IPTG]. GFP was normalized by cell density (OD_600_). For all panels, ANOVA, P < 0.0001, Tukey HSD (* P = 0.0352, ** P ≤ 0.0031, *** P = 0.0002, **** P < 0.0001). For panels C-F, SEM from ≥ 3 biological replicates. Similar trends were observed when GFP was measured at 7 and 9 hours; raw data in Supplementary Fig. 5. **D)** [ATP] for *E. coli* grown in M9 medium with different carbon sources and [IPTG]. For all panels ANOVA, P ≤ 0.0004, Tukey HSD (* P = 0.02, ** P ≤ 0.0079, *** P = 0.0002, **** P < 0.0001). **E)** Growth rate of *E. coli* grown in M9 medium with different carbon sources and [IPTG]. For adenine and ribose, ANOVA, P < 0.0001, Tukey HSD (* P = 0.0108, ** P ≤ 0.0066, **** P < 0.0001). Average residuals for curve fitting in Supplementary Table 3; growth curves in Supplementary Fig. 5. **F)**The effect of changing the carbon source in the growth medium on metabolism normalized by growth. [ATP] and growth rate from panels D and E, respectively

Our quantification of NDM-1 expression, [ATP], and growth rate in medium with glucose provided only a limited parameter space in the large, simulated landscape predicted by our model. To expand the combination of NDM-1 expression, [ATP], and growth rate, we grew bacteria in minimal medium with three additional carbon sources: ribose, adenine, and acetate, which have been shown to alter [ATP] and growth rate (16). Moreover, the carbon sources used were selected to span physiologically relevant metabolic states encountered during infection, including glucose⍰rich conditions with elevated glycolytic flux (40), respiratory growth on short⍰chain carbon sources such as acetate (41), metabolites feeding into the pentose phosphate pathway during infection (42), and host⍰derived purine availability linked to virulence (43). Using the GFP-expressing E. coli strain, we found that increasing [IPTG] consistently increased GFP/OD_600_ across all carbon sources (Fig. 3C), confirming that β-lactamase expression could be systematically tuned across growth environments. However, the magnitude of induction and the significance of the values varied between carbon sources, with ribose producing the largest increase and the greatest range of increases. Similar to our findings with glucose (Fig. 1E), measurements of [ATP] across carbon sources revealed significant, condition-dependent differences (Fig. 3D), and, in nearly all cases, [IPTG] increased [ATP], indicating that NDM-1 expression influences metabolic output. In contrast, growth rate was comparatively insensitive to IPTG and varied primarily with carbon source, with significant changes observed only for adenine and ribose (Fig. 3E). Thus, carbon source and [IPTG] largely influence distinct physiological axes, generating a wide distribution of metabolism normalized by growth values (Fig. 3F) while independently sampling multiple levels of β-lactamase expression.

Using glucose, adenine, ribose, and acetate as carbon sources, we quantified MIC using populations initiated at 10^5^ CFU/mL and 10^4^ CFU/mL (Fig. 4A). Uninduced cultures had a relatively small MIC, which can be attributed to endogenous *ampC* beta-lactamase and/or IE resulting from metabolic and growth dynamics alone (16). We found that increasing [IPTG] in the growth medium consistently and significantly increased ΔMIC relative to the uninduced control, suggesting that IE in our bacteria is mainly due to NDM-1, with *ampC* having a negligible impact. With glucose, ribose, or acetate, increasing [IPTG] significantly increased ΔMIC. However, with adenine, there were no significant differences in ΔMIC amongst all [IPTG] used. We plotted ΔMIC across all carbon sources and [IPTG] as a function of metabolism normalized by growth and observed a biphasic dependence: ΔMIC increased at low values, peaked at intermediate values, and decreased at higher values (Fig. 4B), consistent with model predictions. Neither [ATP] nor growth rate alone predicted ΔMIC, and Gaussian curve fitting showed significantly better agreement when ΔMIC was evaluated as a function of metabolism normalized by growth (Supplementary Fig. 10). ΔMIC showed a significant positive correlation with GFP expression (Fig. 4C), consistent with modeling predictions (Fig. 2A) and the notion that increased β-lactamase expression increases IE (25-29), but insufficient to generate the biphasic trend alone. Importantly, the biphasic relationship was observed only when NDM-1 was expressed. Expression of GFP in the place of NDM-1 yielded a monotonic decrease in ΔMIC with increasing metabolism normalized by growth under GFP⍰inducing conditions (Fig. 4D), as expected in the absence of acquired resistance (16) and consistent with our modeling predictions (Fig. 2E). Furthermore, GFP expression did not correlate with ΔMIC in GFP⍰expressing cells under induced conditions (Fig.□4E), demonstrating that heterologous protein expression alone is insufficient to generate the aforementioned biphasic trend. Together, these results show that while β-lactamase expression contributes to IE, the biphasic dependence of ΔMIC arises from interactions among metabolism, growth, and β-lactamase expression.

**Figure 4:**
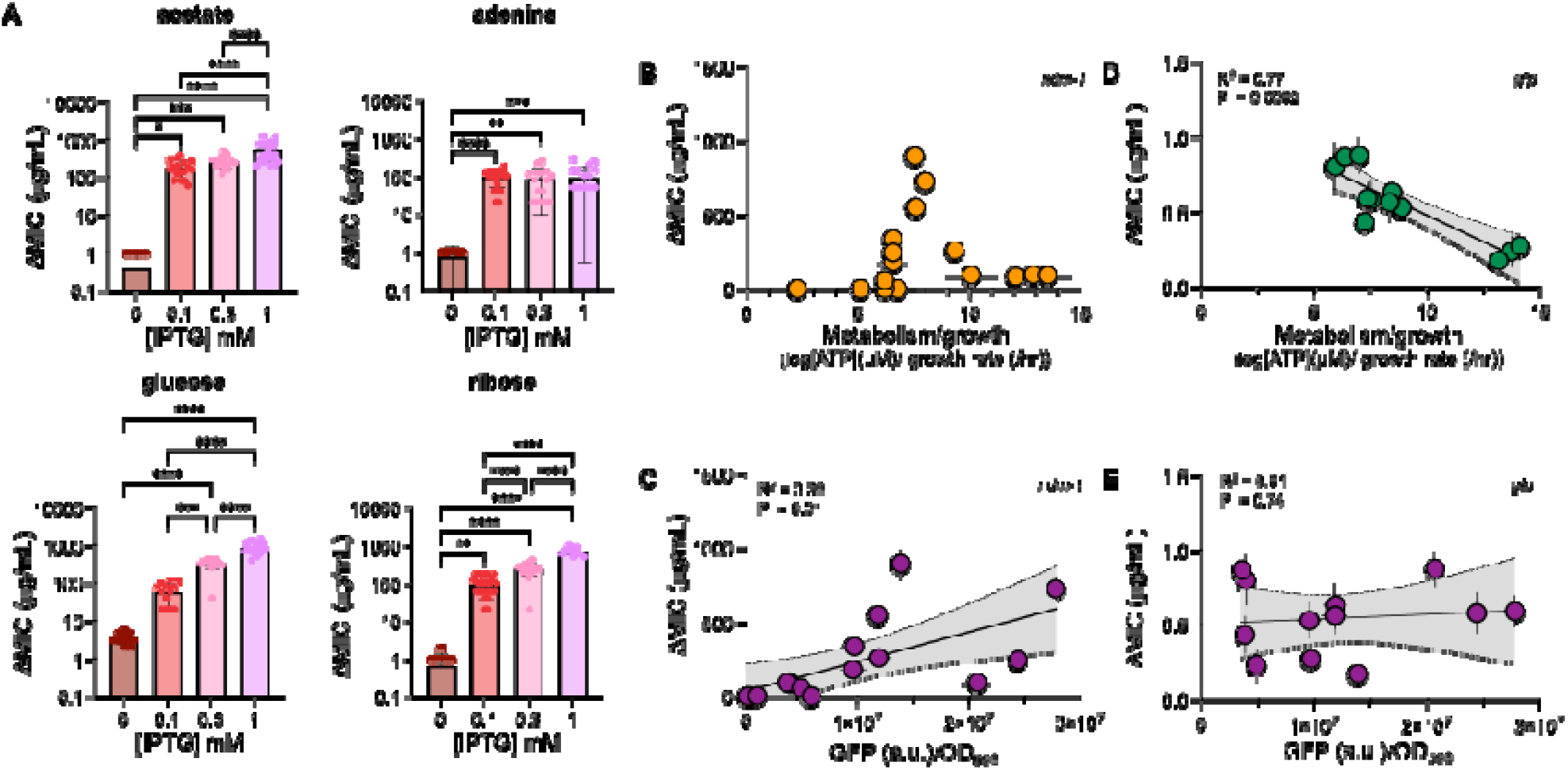
ΔMIC and metabolism normalized by growth follow a biphasic trend when *E. coli* expresses the NDM-1 β-lactamase. A) ΔMIC as a function of [IPTG] for NDM-1 expressing *E. coli* grown in M9 medium with different carbon sources as indicated. SEM from ≥ 4 biological replicates. For all panels, ANOVA, P < 0.0001, Tukey HSD (* P = 0.0196, ** P ≤ 0.0061, *** P ≤ 0.0005,**** P < 0.0001). Raw data in Supplementary Fig. 6 and Fig. 7. B) ΔMIC as a function of metabolism normalized by growth for *E. coli* expressing NDM-1. ΔMIC from panel A; metabolism normalized by growth from Fig. 3F. C) Linear regression of ΔMIC of NDM-1 expressing bacteria as a function of GFP/OD_600_ produced by GFP-expressing bacteria. ΔMIC from panel A; GFP/ OD_600_ from Fig. 3A. Trends were consistent when GFP was measured at 7 and 9 hours (Supplementary Fig. 7) D) ΔMIC as a function of metabolism normalized by growth for *E. coli* expressing GFP. Metabolism normalized by growth in Supplementary Fig. 8. ΔMIC in Supplementary Fig. 8 and 9. For panels D and E, regression with 0 mM IPTG control in Supplementary 10. E) Linear regression of ΔMIC as a function of GFP/OD_600_ for *E. coli* expressing GFP.

### Biphasic dependence of ΔMIC on [ATP]/growth rate is consistent across multiple inocula and β-lactams

Next, we asked whether the biphasic dependence of ΔMIC on metabolism normalized by growth was consistent across multiple initial density comparisons and clinically relevant β-lactam antibiotics. Using cefazolin and imipenem, and using only select combinations of carbon source and [IPTG] to provide a range of metabolism normalized by growth values, we quantified ΔMIC by comparing populations initiated at 10^7^, 10^6^, 10^5^, and 10^4^ CFU/mL (Fig. 5A and D, initial densities for 10^7^ and 10^6^ in Supplementary Fig. 2). Across all density comparisons, where 10^4^ served as the low-density population, and for both antibiotics, ΔMIC exhibited a biphasic dependence on metabolism normalized by growth, with ΔMIC increasing at low values, peaking at intermediate values, and decreasing at higher values (Fig. 5B and E). While the magnitude of ΔMIC varied across initial densities and antibiotics, the overall shape of the relationship was conserved, indicating that the biphasic trend is robust to changes in initial inoculum density and β-lactam type. Consistent with earlier results (Fig. 4D), ΔMIC showed a significant correlation with GFP expression across density comparisons and antibiotics (Fig. 5C and F). Together, these findings indicate that while β-lactamase expression contributes to ΔMIC, the biphasic dependence of ΔMIC on metabolism normalized by growth persists across clinically relevant antibiotics and a broad range of initial bacterial densities.

**Figure 5:**
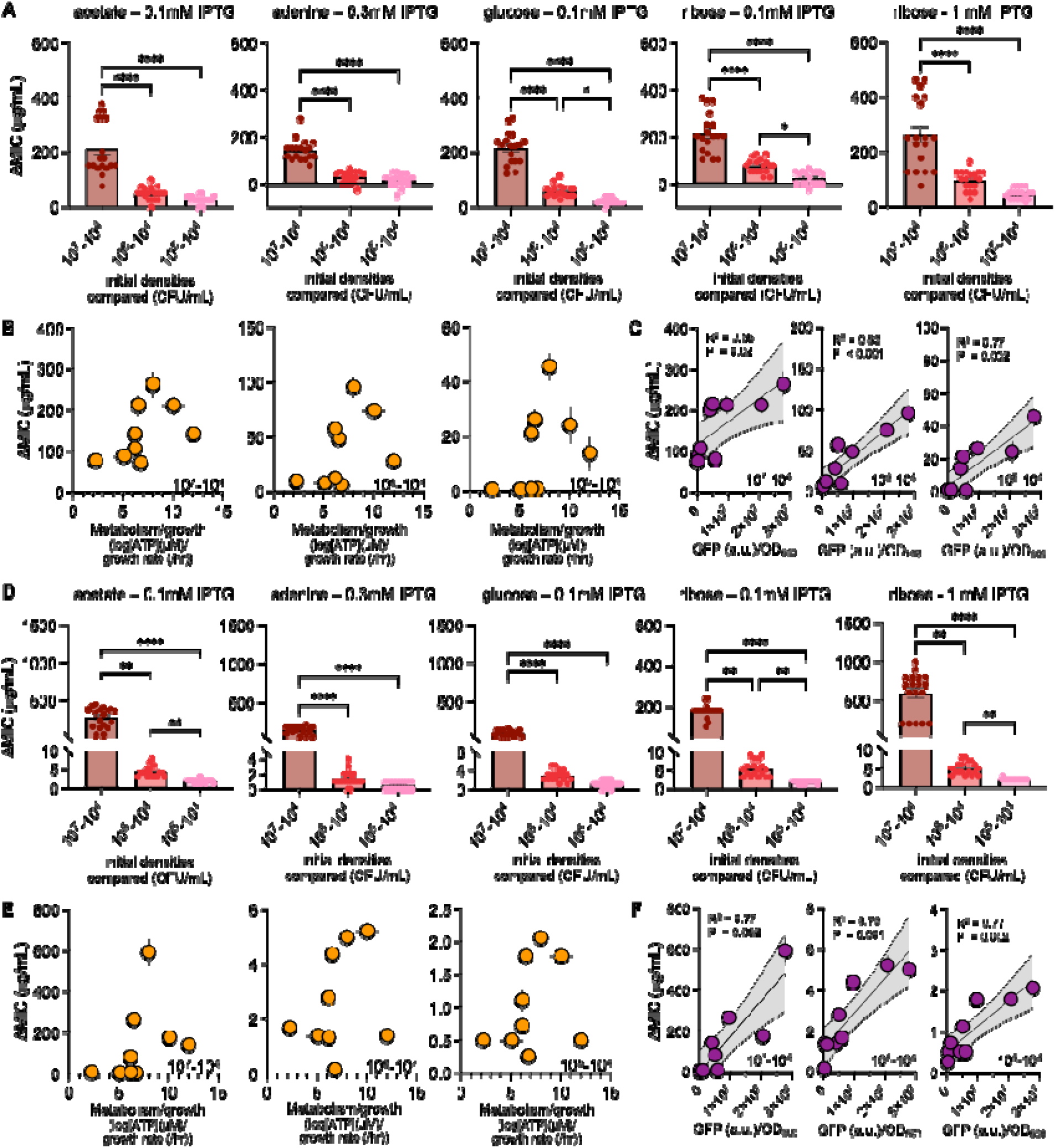
The biphasic relationship between metabolism normalized by growth and ΔMIC is consistent across multiple β-lactam antibiotics and initial cell densities. A) ΔMIC of cefazolin for *E. coli* expressing NDM-1 between different initial densities. Carbon sources and [IPTG] as indicated. SEM from 6 biological replicates. For all panels, Kruskal-Wallis, P < 0.0001, Dunn’s multiple comparison (* P ≤ 0.0301, ** P ≤ 0.0034, *** P = 0.0002, **** P < 0.0001). Raw data in Supplementary Fig. 11. B) ΔMIC of cefazolin as a function of metabolism normalized by growth for *E. coli* expressing NDM-1. Metabolism normalized by growth from Fig. 3F. C) Linear regression of ΔMIC of cefazolin for NDM-1 expressing bacteria as a function of GFP/OD_600_ produced by GFP-expressing bacteria. Initial density comparison as indicated. GFP measured as 5 hours. ΔMIC from panel A; GFP/ OD_600_ from Fig. 3A. D) ΔMIC of imipenem for *E. coli* expressing NDM-1 between different initial densities. Carbon sources and [IPTG] as indicated. SEM from 6 biological replicates. For all panels, Kruskal-Wallis, P < 0.0001, Dunn’s multiple comparison (* P = 0.0329, ** P = 0.0034, *** P = 0.0008, **** P < 0.0001). Raw data in Supplementary Fig. 12. E) ΔMIC of imipenem as a function of metabolism normalized by growth for *E. coli* expressing NDM-1. Metabolism normalized by growth from Fig. 3F. F) Linear regression of ΔMIC of imipenem for NDM-1 expressing bacteria as a function of GFP/OD_600_ produced by GFP-expressing bacteria. Initial density comparison as indicated. GFP measured as 5 hours. ΔMIC from panel A; GFP/ OD_600_ from Fig. 3A.

### The biphasic relationship persists in clinical isolates of β-lactamase-expressing E. coli

To determine whether the biphasic dependence of ΔMIC on metabolism normalized by growth observed in our reductionist system extends to clinical isolates, we selected *E. coli* strains from a Gram-negative isolate bank encoding β-lactamases that either confer resistance to the clinically relevant carbapenem imipenem (NDM- and KPC-type) or do not (TEM-, CMY-, and EC-type). When grown in M9 medium with glucose as the carbon source, these isolates exhibited significant differences in [ATP] per cell (Fig. 6A) and in growth rate (Fig. 6B), resulting in a broad range of metabolism normalized by growth values across the clinical panel (Fig. 6C). We next quantified ΔMIC using average initial densities of 10^7^ and 10^5^ CFU/mL, which are commonly used to define IE in clinical susceptibility testing (44). Using a CSLI-like gradient of imipenem, isolates expressing NDM- or KPC-type β-lactamases exhibited high ΔMIC values (Fig. 6D). In contrast, isolates without NDM or KPC-type, but instead containing TEM-, CMY-, or EC-type β-lactamases showed uniformly low ΔMIC (Fig. 6E). When ΔMIC was plotted as a function of metabolism normalized by growth, NDM- and KPC-expressing isolates exhibited a biphasic dependence (Fig. 6F). In contrast, analysis of all non-resistant β-lactamase–expressing isolates together yielded a weak, non-significant association between ΔMIC and metabolism normalized by growth (Fig. 6G). However, separating these isolates by β-lactamase class revealed stronger linear relationships that approached statistical significance (Fig. 6H), likely reflecting the limited number of isolates containing each β-lactamase type. Our model predicts this behavior, with simulations reproducing the observed negative relationship between ΔMIC and metabolism normalized by growth (Fig.□6H). This trend also parallels that of GFP⍰expressing cells, where heterologous protein expression alone does not confer antibiotic protection, and ΔMIC declines with increasing metabolism normalized by growth (Fig.□4D) (16). Together, these results demonstrate that the biphasic relationship between ΔMIC and metabolism normalized by growth occurs in clinical *E. coli* isolates and that its presence and form depend on the specific β-lactamase–β-lactam pair, establishing physiological relevance without predicting patient-level outcomes.

**Figure 6:**
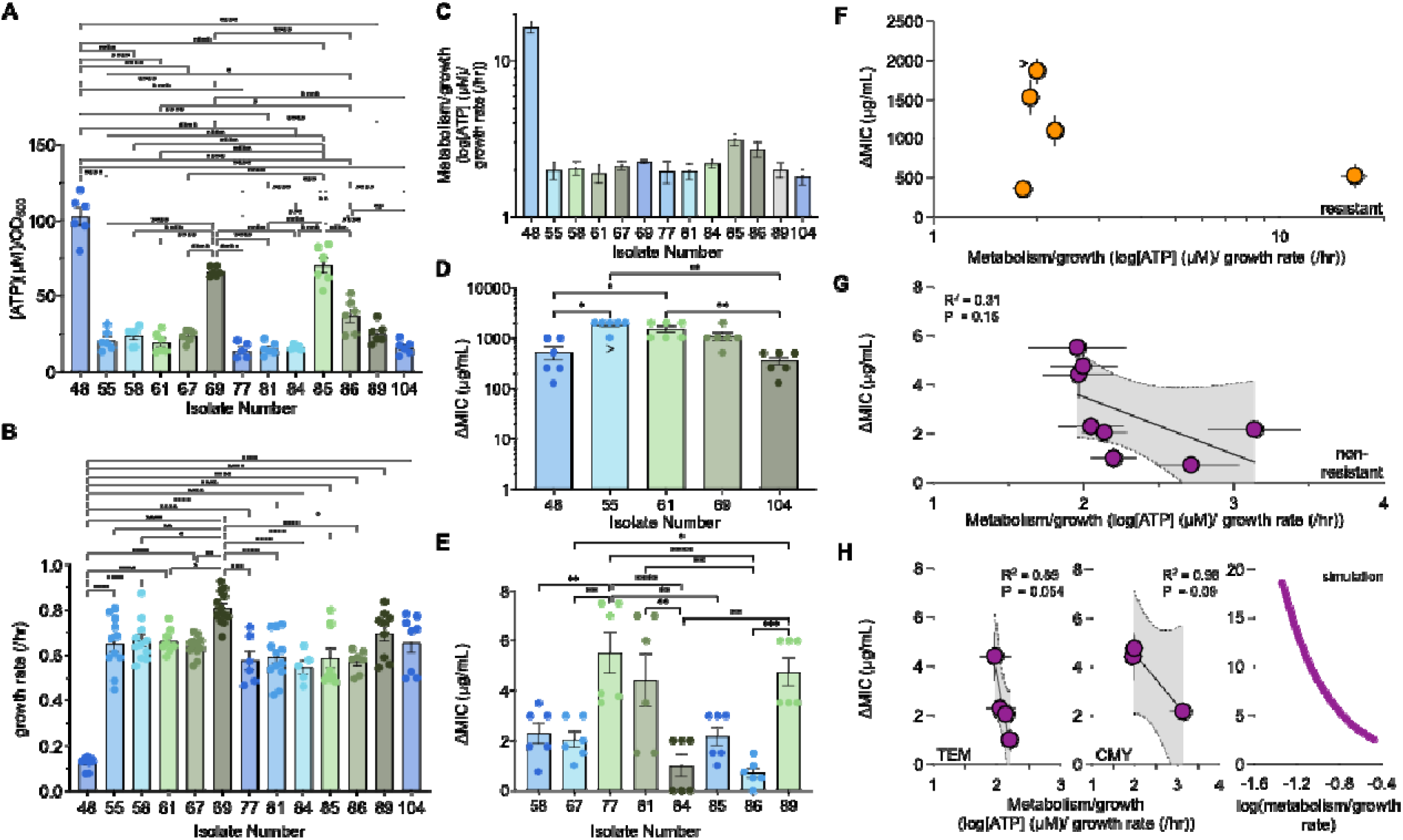
The relationship between ΔMIC and metabolism normalized by growth depends on β-lactamase type in clinical isolates of *E. coli*. A) [ATP] for *E. coli* clinical strains grown in M9 medium with glucose and 0.1% CAA. SEM from ≥ 5 biological replicates. ANOVA, P ≤ 0.0001, Tukey HSD (* P ≤ 0.0282, ** P ≤ 0.0021, *** P = 0.0005, **** P < 0.0001). Strain number is listed on the x-axis. Strain information in Supplementary Table 5. B) Growth rate for *E. coli* clinical strains grown in M9 medium with glucose and 0.1% CAA. SEM from ≥ 5 biological replicates. ANOVA, P ≤ 0.0001, Tukey HSD (* P ≤ 0.0489, ** P ≤ 0.0076, *** P = 0.0005, **** P < 0.0001). Raw growth curves in Supplementary Fig. 13. Residuals for curve fitting in Supplementary Table 6. C) Metabolism normalized by growth *E. coli* clinical strains. Data from panels A and B. Error bars = SEM. D) ΔMIC of imipenem for *E. coli* clinical strains expressing either NDM-1 or KPC type β-lactamases. SEM from 6 biological replicates. Kruskal-Wallis, P = 0.0003, Dunn’s multiple comparison test (* P ≤ 0.0198, ** P ≤ 0.0083). Raw data in Supplementary Fig 14. For isolate 55, (panels D and E) > indicates that ΔMIC is greater than the value reported, as some replicates showed growth at their highest concentration of imipenem that could be tested. E) ΔMIC of imipenem for *E. coli* clinical strains expressing TEM, CMY, or EC type β-lactamases. SEM from 6 biological replicates. ANOVA, P < 0.0001, Tukey HSD (* P = 0.0440, ** P ≤ 0.0084, *** P = 0.0005, **** P < 0.0001). F) ΔMIC of imipenem as a function of metabolism normalized by growth for *E. coli* clinical strains expressing NDM-1 or KPC-type β-lactamase. Metabolism normalized by growth from panel C. G) Linear regression of ΔMIC of imipenem as a function of metabolism normalized by growth for *E. coli* clinical strains expressing TEM, CMY, or EC type β-lactamase. Metabolism normalized by growth from panel C. H) Left: Linear regression of ΔMIC of imipenem as a function of metabolism normalized by growth for *E. coli* clinical strains expressing TEM-type β-lactamases. Center: Linear regression of ΔMIC of imipenem as a function of log(ATP)/growth rate for *E. coli* clinical strains expressing CMY-type β-lactamases. Right: Simulations showing the absence of β-lactamase expression (β=0) in our model.

## Discussion

We have shown that expressing a β-lactamase can non-intuitively alter [ATP] while often leaving the growth rate unchanged, thereby altering metabolism normalized by growth. Using diverse carbon sources, we found that ΔMIC exhibits a biphasic dependence on metabolism normalized by growth, with intermediate values yielding the greatest IE strength. Simulations suggest that this is due to interactions between metabolism, growth rate, and β-lactamase expression. This trend is β-lactamase-specific, as GFP expression abolished the biphasic relationship between metabolism normalized by growth. Importantly, this biphasic dependence was observed not only in a reductionist laboratory system but also in clinical *E. coli* isolates, demonstrating that this relationship extends to clinically relevant situations. Together, our results identify a simple rule: IE in β-lactamase-expressing bacteria is maximized at intermediate values of metabolism relative to growth. Below this regime, resistance⍰mediated protection dominates antibiotic outcomes; above it, metabolism⍰potentiated lethality overwhelms resistance. While we focus on β⍰lactams, the underlying competition between resistance⍰mediated protection and physiology⍰dependent lethality is likely relevant to other resistance–antibiotic pairs.

We found that increasing NDM-1 expression with IPTG generally increased [ATP] but had only carbon-source-specific effects on growth rate. While heterologous protein expression is often assumed to impose a metabolic burden that slows growth (23, 24, 45), our results demonstrate that this is not always the case. Indeed, acquisition of some, but not all, β-lactamases can increase growth rate even in the absence of antibiotics (23). To our knowledge, we are the first to demonstrate an increase in [ATP] resulting from β-lactamase expression. Increases in [ATP] following heterologous protein expression have been reported previously (46) and may reflect compensatory changes in cellular respiration. We caution that [ATP] should be interpreted as a proxy for broader metabolic state, correlating with metrics such as the NAD^+^/NADH ratio and oxygen consumption (14), rather than as a singular causal mediator or indicator of specific metabolic pathways driving antibiotic susceptibility

As predicted by our mathematical model (Fig.□2) and supported by our experiments, the observed biphasic relationship between ΔMIC and metabolism normalized by growth shows that IE is maximized at intermediate values of this metric. This phenomenon can be understood by first considering model predictions for how metabolism normalized by growth determines ΔMIC in the absence of β⍰lactamase. β⍰lactamase expression then reshapes, rather than replaces, this underlying physiological relationship. As previously described (16), low-density populations spend more time in metabolically active log-phase growth (Fig. 7). They are therefore more susceptible to metabolism-potentiated antibiotic lethality (14), whereas high-density populations rapidly enter the stationary phase and are relatively protected. Growth rate determines how quickly these populations exit log phase, while [ATP] reflects the intensity of metabolism during this period. Environments with fast growth relative to low [ATP] exacerbate IE, whereas slow growth relative to high [ATP] minimizes it (16). These previously reported relationships in bacteria lacking β-lactamases show a monotonic, negative linear relationship between ΔMIC and metabolism normalized by growth (Fig. 7), whether cultures are initiated directly from overnight cultures or in mid-log phase (16). Importantly, our data confirm these model predictions in the absence of a β-lactamase (Fig. 4D) and show that β-lactamase activity reshapes, but does not override, this physiological framework.

**Figure 7:**
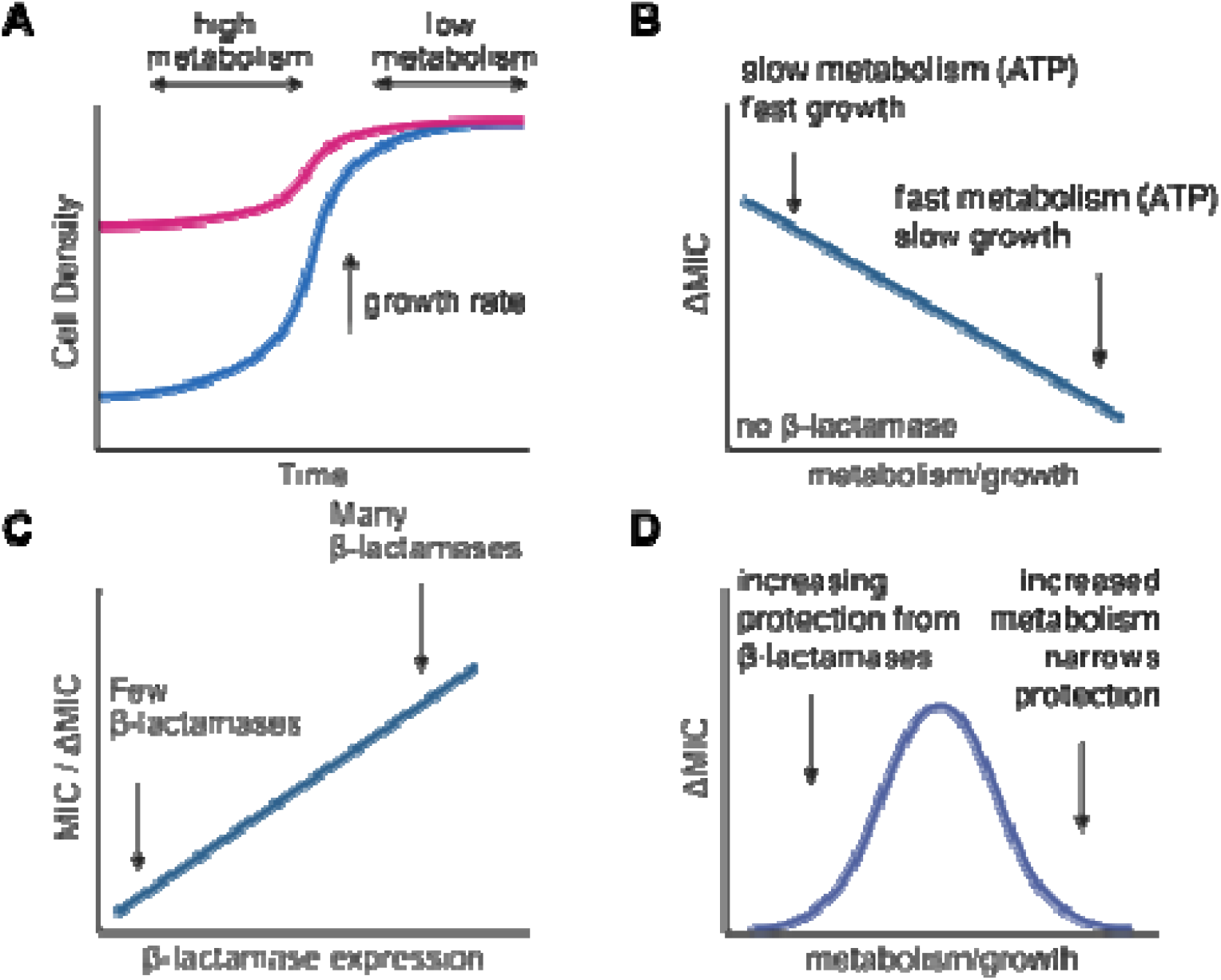
Our proposed mechanism to explain the biphasic dependence of ΔMIC on metabolism normalized by growth. A) The initial density of a bacterial population determines how long it spends in log phase growth, where metabolism is greater. The growth rate afforded to the population dictates how long it takes for the population to reach the stationary phase, where metabolism is reduced. B) When growth is relatively greater than metabolism (e.g., ATP), the difference in MIC between both populations is increased as the high-density population spends a relatively shorter time in log phase growth, where antibiotic lethality is maximized. This increases ΔMIC. When [ATP] is greater than growth, the time at which both populations have high metabolism becomes increasingly similar, leading to similar MICs between them. This reduces ΔMIC. C) Independent of changes in metabolism and growth, increasing β-lactamase expression increases ΔMIC; the more β-lactamases available, the more the population collectively inactivates β-lactams. D) The biphasic relationship between ΔMIC and metabolism normalized by growth observed in our study. On the left side of the curve, increasing β-lactamase expression increases ΔMIC. Owing to low metabolism relative to growth, the difference in ΔMIC between high and low density populations is increased. On the right side of the curve, increases in metabolism relative to growth allow both the high- and low-density populations to inactivate similar amounts of β-lactam antibiotics, reducing the difference in MIC and leading to lower ΔMIC.

Expressing a β-lactamase partially alters this relationship by increasing ΔMIC through antibiotic degradation, consistent with model predictions (Fig.□2) and thereby explaining the left side of the biphasic trend (Fig. 7). At low metabolism normalized by growth, increased β-lactamase expression accentuates the difference in MIC between high- and low-density populations. However, at higher metabolism normalized by growth, increased metabolism potentiates antibiotic lethality such that both populations experience similar levels of killing despite β-lactamase expression, reducing ΔMIC. Consistent with this interpretation, a biphasic relationship was not observed when β-lactamase expression or metabolic state varied alone (Supplementary Fig. 4). Our GFP controls indicate that heterologous protein expression alone is insufficient to generate this biphasic trend. We cannot rule out reduced expression or dysfunctional β-lactamase activity at high metabolism normalized by growth, which may also explain the rightward slope of the biphasic trend, although we found no relationship between GFP expression and metabolism normalized by growth (Supplementary Fig. 10). Importantly, differences in carbon source alone cannot explain the biphasic trend, as multiple carbon sources are represented on the left and right of the curve (Fig. 4B).

Analysis of clinical isolates further revealed that the emergence of a biphasic relationship depends on both the β-lactamase class and the β-lactam antibiotic. Isolates expressing NDM- or KPC-type β-lactamases exhibited biphasic behavior with imipenem, whereas isolates expressing TEM- or CMY-type enzymes showed uniformly low ΔMIC and a negative relationship with metabolism normalized by growth. This antibiotic-specific dependence likely reflects differences in enzymatic activity and susceptibility profiles, underscoring that biphasic IE behavior is not universal across all β-lactamase–β-lactam combinations. While these clinical isolate experiments do not directly predict patient-level outcomes, they demonstrate that biphasic ΔMIC–metabolism relationships emerge within clinically relevant isolates.

Our results show that the carbon source in the growth medium shapes β-lactamase-mediated IE, highlighting that nutrient availability, rather than β-lactamase expression alone, critically influences IE strength. This finding is consistent with previous work showing that the composition of the growth environment can strongly influence antibiotic efficacy, often by altering bacterial metabolism to potentiate or reduce antibiotic lethality (47, 48) . Accordingly, we found that changes in carbon source, together with NDM-1 expression, altered bacterial metabolism. This dependence of ΔMIC on the composition of the growth environment may help explain why some clinical studies have failed to detect IE (7, 49), whereas others report pronounced effects on patient outcomes, including mortality (50, 51). It further suggests that different infection sites may vary in their propensity to exhibit IE, as nutrient availability differs across host environments (52, 53), and helps explain why the presence of a β-lactamase alone cannot reliably predict IE occurrence, (30, 31) as IE may only emerge under specific growth environments. Defining growth environments in which IE is amplified or suppressed may therefore improve predictions of antibiotic efficacy *in vivo*.

## Methods

### Bacterial strains, growth conditions, and antibiotics

*E. coli* strain DH5αPRO (Clontech, Mountain View, CA) was used for our reductionist system. *E. coli* expressing *bla*_*NDM-1*_ were created as described previously (54). *bla*_*NDM-1*_ was amplified from the plasmid pGDP1 NDM-1 plasmid (Addgene, Watertown, MA) and cloned into plasmid pPROLAR (origin of replication = p15a, selectable marker = kanamycin) using EcoR1 (New England Biolabs, Ipswich, MA) and HindIII (New England Biolabs). The resulting plasmid was sequenced and transformed into DH5αPRO. As a control, *gfpmut(2b*) was cloned into pPROLAR using EcoR1 and HindIII. Thus, plasmid maintenance effects are inherently incorporated through the matched plasmid backbone and GFP controls used throughout the study. *E. coli* clinical isolates were obtained from the Gram-Negative Carbapenemase Detection (CarbaNP) provided by the AR Isolate Bank at the Centers for Disease Control.

To create overnight cultures from which each experiment was initiated, single colonies were isolated from the lysogeny broth (LB) (MP Biomedical Santa Ana, CA) agar and inoculated into 3 mL of LB liquid medium with 50 *μ*g/mL of kanamycin sulphate (ThermoFisher Scientific, Pittsburgh, PA) contained in 15 mL culture tubes (Genesee Scientific, Morrisville, NC). Overnight cultures were created by shaking the culture tubes at 37°C and 250 revolutions per minute for 24 hours.

All experiments were conducted in modified M9 medium [1X M9 salts (48 mM Na_2_HPO_4_, 22 mM KH_2_PO_4_, 862 mM NaCl, 19 mM NH_4_Cl), 0.5% thiamine (Alfa Aesar, Ward Hill, MA), 2 mM MgSO_4_, 0.1 mM CaCl_2_], supplemented with 0.04% of each carbon source. For our reductionist system, media contained varying concentrations of casamino acids (0.1%, 0.5% and 1%, Teknova, Hollister, CA) and isopropyl β-D-1-thiogalactopyranoside (IPTG, ThermoFisher). The carbon sources tested in this study were sodium acetate (Sigma-Aldrich, St. Louis, MO), adenine (Thermo Fisher Scientific), D-glucose (Thermo Fisher Scientific), and D-ribose (Thermo Fisher Scientific). For experiments with clinical isolates, we used modified M9 medium with glucose and 0.1% CAA. Ampicillin sodium salt (ThermoFisher), cefazolin sodium salt (ThermoFisher), and imipenem monohydrate (MedChemExpress, Monmouth Junction, NJ) were used to challenge *E. coli*.

### Measuring ATP

We measured [ATP] using a previously published protocol that separately measures [ATP] from changes in growth (14). Overnight cultures were diluted 1/40 into 40 mL of M9 medium in a 50 mL conical tube with the lid slightly unscrewed to allow gas exchange. The M9 medium contained IPTG but lacked casamino acids and antibiotics. Bacteria were shaken for 2 hours at 250 RPM/37^°^C. Cultures were concentrated at least 2X (OD_600_ of > 0.05) in diluted (3:1) M9 medium that contained a carbon source (0.04%), IPTG (0, 0.1, 0.3, or 1mM), and casamino acids (0.1%, 0.5%, or 1%). 100 μL of these cultures were added to the wells of an opaque-walled 96-well plate (Costar 3370, Corning, Kennebunk, ME) in technical duplicate. The plate was covered with 2 BreathEasy (Sigma-Aldrich) membranes and placed in a shaker for 1 hour at 250 RPM/37^°^C. [ATP] was then measured using a BacterTiter-Glo microbial cell viability assay (Promega, Madison, WI) according to manufacturer recommendations and using a VICTOR X4 microplate reader (Perkin Elmer, Waltham, MA). Luminescence values were normalized to optical density (OD) at 600 nm (OD_600_). [ATP] was determined using a standard curve produced using purified ATP (16). Similar trends are observed when ATP is measured at 3 and 5 hours post-addition of casamino acids (16).

### Growth rate

Overnight cultures were resuspended in dH_2_O. Each culture was diluted 200-fold into 200 μL of M9 medium (containing a carbon source, casamino acids, and IPTG) that was housed in a 96-well plate. The wells were overlaid with 70 μL of mineral oil to prevent evaporation. Cell density (OD_600_) was measured every 10 minutes for 10 hours in a Victor X4 (Perkin Elmer) microplate reader preset to 37^°^C. OD_600_ values from cell-free medium were subtracted from all measurements. The maximum growth rate was determined from the experimental growth curves by fitting the data to a logistic growth equation as previously described.

### MIC assays – reductionist system and gfp-expressing strain

Overnight cultures of the reductionist system were resuspended in dH_2_O and diluted to either 10^7^ (2.87 x 10^7^ ± 5.36 x 10^6^ CFU/mL), 10^6^ (2.70 x 10^6^ ± 1.00 x 10^5^ CFU/mL), 10^5^ (5.3 x 10^5^ ± 1.24 x 10^4^ CFU/mL) or 10^4^ (2.34 x 10^4^ ± 3.16 x 10^3^ CFU/mL, Supplementary Fig. 2) into 200 μL of M9 medium housed in a 96-well plate (non-treated, Genesee Scientific) containing increasing concentrations of antibiotic and IPTG. The GFP expressing strain was treated similarity with initial densities of 10^7^ (2.30 x 10^7^ ± 6.66 x 10^6^ CFU/mL), 10^6^ (2.13 x 10^6^ ± 5.81 x 10^5^ CFU/mL), 10^5^ (1.67 x 10^5^ ± 4.26 x 10^4^ CFU/mL) or 10^4^ (2.57 x 10^4^ ± 6.57 x 10^3^ CFU/mL, Supplementary Fig. 2). The plates were sealed with two BreathEasy membranes (Sigma-Aldrich) and were placed in a shaker for 24 hours at 250 RPM and 37°C. After 24 hours, cell density (OD_600_) was measured using a Victor NIVO plate reader (Perkin Elmer). Measurements below 0.01 were set to zero, as values below this threshold are not reliable indicators of growth in our system (16). Our MIC assay has previously been shown to yield consistent results when OD_600_ was measured at 24 and 40 hours and when CFUs are enumerated at 24 hours (16). MIC was defined as the lowest antibiotic concentration that inhibited growth. To assess the strength of IE (ΔMIC), we used the following equation:

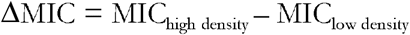

Initial density was measured from overnight cultures using a dilution series followed by plating on LB agar containing 50 *μ*g/mL of kanamycin. CFUs were enumerated after 20 hours of incubation at 37 °C.

### MIC assays – clinical isolates

Overnight cultures were resuspended in dH_2_O and diluted to either 10^7^ (average of all strains; 2.80 x 10^7^ ± 2.39 x 10^6^ CFU/mL) or 10^5^ (average of all strains; 3.46 x 10^5^ ± 2.56 x 10^4^ CFU/mL, Supplementary Fig. 13) into 200 *μ*L of M9 medium (glucose, 0.1% CAA) housed in a 96-well plate (non-treated, Genesee Scientific) containing increasing concentrations of imipenem monohydrate. Mimicking CSLI standards, [imipenem] was started at 0.125 µg/mL and increased 2-fold to 2048 µg/mL, the highest concentration achievable due to solubility limits. Plates were then grown and quantified as above. ΔMIC was quantified as above. Initial density was measured as above without kanamycin selection.

### GFP expression

To measure the effect of increasing IPTG on GFP expression, GFP-expressing *E. coli* were grown overnight, washed once with dH_2_O, and diluted 100-fold into 200 μL of M9 medium in a 96-well microplate (non-treated, Genesee Scientific). The microplate was covered with two BreathEasy films (Sigma-Aldrich), and shaken at 37 °C/250 RPM. After 3, 5, 7, and 9 hours, GFP and OD_600_ were measured in a Victor NIVO microplate reader. To process the data, GFP and OD_600_ from the cell-free media were first subtracted from all samples. GFP was then normalized by OD_600_. Finally, the GFP signal from uninduced cultures was removed from all measurements. Any negative measurements were set to zero.

To determine differences in GFP production rate as a function of initial population density, GFP-expressing bacteria were grown as above and then diluted to 10^7^, 10^6^, 10^5,^ or 10^4^ in M9 medium in a 96-well microplate (non-treated, Genesee Scientific). The microplate was covered with two BreathEasy films (Sigma-Aldrich) and shaken at 37 °C/250 RPM. GFP and OD_600_ were measured in a Victor NIVO during mid-log phase for each population as determined by trends in OD_600_. GFP signal was processed as above. GFP expression rate was determined by fitting linear lines through plots of GFP vs OD_600_ over time (hours).

### Mathematical modeling

We modified a previously published model to capture IE in the expression of β-lactamase (16).

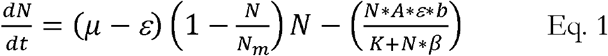

In our model, bacterial growth is captured by a logistic growth equation (on the left). Bacterial growth depends upon two parameters: the growth rate (*μ*,/hr) of the bacteria and non-biomass-generating metabolism (*ε*, mmol/g/hr), which approximates a maintenance coefficient. Bacteria (*N*, unitless) are initiated from one of two initial densities, high *(N*_*high*_) or low (*N*_*low*_), and grow to a fixed carrying capacity (*N*_*m*_, unitless). Antibiotics reduce logistic growth via a *per capita* death rate term (on the right), modeled after Michaelis-Menten dynamics. The value of this term is dependent upon the bacterial density (*N*), the concentration of antibiotics (*A*, unitless), the half maximal rate of antibiotic-induced death (*K*, unitless), antibiotic lethality (*b*, /*hr*), metabolism (*ε*, mmol/g/hr), and the expression of β-lactamases (*β*, unitless). To account for the collective degradation of antibiotics by β-lactamases, we scaled *β* by N; the greater the bacterial density, the more β-lactamases are produced (25). This serves to reduce the impact of antibiotic lethality. To capture the metabolism-inducing increase of antibiotic lethality (14), *A* is scaled by *ε*. All simulations were performed for *t* = 24 hours using MATLAB (R2019b, MathWorks Inc., Natick, MA) using ode45.

We estimated the maximum growth rate (*μ*) using the average growth rate of *E. coli* in M9 medium (0.301 ± 0.003/hr) from our experiments. We used previously reported maintenance coefficients (0.013-0.473 mmol/g/hr(55)) to estimate the order of magnitude of metabolism. To determine the range over which β-lactamase expression (*β*) could vary, we considered previously produced induction curves showing ∼250-fold increase in luciferase activity as [IPTG] is increased from 0 mM to 1 mM and when driven by the P_lac-ara-1_ promoter (35). When *β* = 1, killing kinetics of the right-hand side of Eq. 1 are solely driven by the half maximal killing rate of the antibiotics (*K*) and cell density (*N*) in the denominator, capturing IE without an active form of resistance(16). Thus, *β* values greater than 1 are required for β-lactamase expression in our model. Previously published time-kill curves showing that 0.1%-1% of bacteria survive after 3 hours in antibiotics provided at a concentration greater than MIC (14) were used to estimate the antibiotic killing rate (*b*). To capture the general trend of this range, we assumed a value of 0.1. The half-maximal killing rate of the antibiotic (*K*) was estimated by fitting the simulation results to the qualitative trends of our data. We estimated the initial density of the 10^5^ and 10^4^ CFU/mL populations by assuming that the carrying capacity of M9 medium with 0.04% glucose and 0.1% casamino acids was ∼10^8^ CFU/mL. Experimentally, the initial density of high and low populations in our study was determined to be 5.3 x 10^5^ ± 1.24 x 10^4^ CFU/mL and 2.34 x 10^4^ ± 3.16 x 10^3^ □CFU/mL, respectively. When normalized to a carrying capacity of 10^8^ CFU/mL, this corresponded to initial densities ranging from 10^-2^ to 10^-3^ and from 10^-3^ to 10^-4^ for high and low density, respectively. We therefore set our initial densities to 5 x 10^-2^ (*N*_*high*_) and 1 x 10^-4^ (*N*_*low*_) in our model. Sensitivity analysis for key model parameters is presented in Supplementary Fig. 4. All core parameters are listed in Supplementary Table 2; modifications to this parameter set are described throughout.

### Statistical analysis

GraphPad Prism (version 11, San Diego, California) was used to test for statistical significance throughout, except for Gaussian curve fitting, which was performed using JMP (version 18.2.2, SAS Institute, Cary, NC). Data distributions were assessed for normality using the Shapiro– Wilk test, and appropriate parametric or non-parametric statistical tests were applied as indicated in the figure legends and text. [ATP] and growth rate measurements used to calculate log[ATP]/growth rate (metabolism normalized by growth) were obtained from independent experiments that were not matched in biological replicates. As a result, [ATP] and growth rate values could not be paired at the individual-replicate level, precluding formal statistical testing of the derived metabolism normalized by growth metric. Instead, metabolism normalized by growth was calculated using the mean [ATP] and mean growth rate values pooled across all independent replicates for a given condition. The standard error of the mean (SEM) for metabolism normalized by growth was estimated by propagating the errors from the SEMs of [ATP] and growth rate measurements. As a result, derived metabolism normalized by growth values is presented as a descriptive metric to compare trends across conditions. ATP was log-transformed prior to normalization by growth rate to stabilize scale differences arising from order-of-magnitude variation across conditions.

## Author contributions

Conceptualization - ARKT and RPS

Formal analysis –ARKT, MBM, and RPS

Funding acquisition – RPS and AJL

Investigation – ARKT, KS, PI, TM, EMM, YN, DPM, HV, NN, JG, TR, DPM, DW, TW, AP, RPS

Methodology - ARKT

Project administration – RPS

Supervision – RPS

Visualization – ARKT, RPS

Writing – original draft - ARKT, RPS

Writing – review & editing - ARKT, KS, PI, TM, EMM, YN, DPM, HV, NN, JG, TR, DPM, DW, TW, AP, AJL

## Acknowledgements

This work was supported by National Institutes of Health grant R15AI159902 (RPS) and National Institutes of Health grant 1R35GM150871-01 (AJL). Images were created using Biorender.

## Data availability

All data associated with this manuscript can be found on figshare at the following address: https://figshare.com/s/3d0cd955a9bda863cec8

## Code availability

The modeling code for our mathematical model is available on figshare at the following address: https://figshare.com/s/3d0cd955a9bda863cec8. Modeling code for curve fitting is available in the Dryad Digital Repository associated with (16).

## References

1. I. Brook, Inoculum effect. Reviews of infectious diseases 11, 361–368 (1989).

2. C. Tan et al., The inoculum effect and band-pass bacterial response to periodic antibiotic treatment. Molecular Systems Biology 8, 617 (2012).

3. J. R. Lenhard, Z. P. Bulman, Inoculum effect of β-lactam antibiotics. Journal of Antimicrobial Chemotherapy 74, 2825–2843 (2019).

4. K. P. Smith, J. E. Kirby, The inoculum effect in the era of multidrug resistance: minor differences in inoculum have dramatic effect on MIC determination. Antimicrobial agents and chemotherapy 62, 10.1128/aac.00433–00418 (2018).

5. B. Jean et al., β-Lactam Inoculum Effect in Methicillin-Susceptible Staphylococcus aureus Infective Endocarditis. JAMA Netw Open 7, e2451353 (2024).

6. W. R. Miller et al., The Cefazolin Inoculum Effect Is Associated With Increased Mortality in Methicillin-Susceptible Staphylococcus aureus Bacteremia. Open Forum Infectious Diseases 5, ofy123 (2018).

7. S. Lee et al., Clinical implications of cefazolin inoculum effect and β-lactamase type on methicillin-susceptible Staphylococcus aureus bacteremia. Microbial Drug Resistance 20, 568–574 (2014).

8. J. Martinez, F. Baquero, Mutation frequencies and antibiotic resistance. Antimicrob Agents Chemother 44, 1771–1777 (2000).

9. B. Fantin et al., The inoculum effect of Escherichia coli expressing mcr-1 or not on colistin activity in a murine model of peritonitis. Clinical Microbiology and Infection 25, 1563. e1565.–1563.e1568 (2019).

10. H. Eagle, The effect of the size of the inoculum and the age of the infection on the curative does of penicillin in experimental infections with Streptococci, Pneumococci and Treponema pallidum. The Journal of Experimental Medicine 90, 595–607 (1949).

11. M. Ahmad, S. V. Aduru, R. P. Smith, Z. Zhao, A. J. Lopatkin, The role of bacterial metabolism in antimicrobial resistance. Nature Reviews Microbiology, 1–16 (2025).

12. E. Tuomanen, R. Cozens, W. Tosch, O. Zak, A. Tomasz, The rate of killing of Escherichia coli by beta-lactam antibiotics is strictly proportional to the rate of bacterial growth. J Gen Microbiol 132, 1297–1304 (1986).

13. U. Łapińska et al., Fast bacterial growth reduces antibiotic accumulation and efficacy. Elife 11, e74062 (2022).

14. A. J. Lopatkin et al., Bacterial metabolic state more accurately predicts antibiotic lethality than growth rate. Nature microbiology 4, 2109–2117 (2019).

15. D. M. Hernandez et al., Purine and pyrimidine synthesis differently affect the strength of the inoculum effect for aminoglycoside and B-lactam antibiotics. bioRxiv, 2024.2004. 2009.588696 (2024).

16. G. Diaz-Tang et al., Growth productivity as a determinant of the inoculum effect for bactericidal antibiotics. Science Advances 8, eadd0924 (2022).

17. C. L. Tooke et al., β-Lactamases and β-Lactamase Inhibitors in the 21st Century. Journal of molecular biology 431, 3472–3500 (2019).

18. M. A. Wikler, Performance standards for antimicrobial susceptibility testing Sixteenth informational supplement. M 100-S 16 (2006).

19. J. C. McNeil et al., Cefazolin inoculum effect and methicillin-susceptible Staphylococcus aureus osteoarticular infections in children. Antimicrob Agents Chemother 64, e00703–00720 (2020).

20. C. Betriu et al., Comparative in vitro activity and the inoculum effect of ertapenem against Enterobacteriaceae resistant to extended-spectrum cephalosporins. Int J Antimicrob Agents 28, 1–5 (2006).

21. Y.-A. Na et al., Growth retardation of Escherichia coli by artificial increase of intracellular ATP. Journal of Industrial Microbiology and Biotechnology 42, 915–924 (2015).

22. G. Wu et al., Metabolic burden: cornerstones in synthetic biology and metabolic engineering applications. Trends in biotechnology 34, 652–664 (2016).

23. F. Rajer, L. Sandegren, The role of antibiotic resistance genes in the fitness cost of multiresistance plasmids. mBio 13: e0355221. DOI 10,03552–03521 (2022).

24. D. C. Marciano, O. Y. Karkouti, T. Palzkill, A fitness cost associated with the antibiotic resistance enzyme SME-1 β-lactamase. Genetics 176, 2381–2392 (2007).

25. R. P. Smith et al., The public and private benefit of an impure public good determines the sensitivity of bacteria to population collapse in a snowdrift game. Environmental microbiology 21, 4330–4342 (2019).

26. K. S. Thomson, E. S. Moland, Cefepime, piperacillin-tazobactam, and the inoculum effect in tests with extended-spectrum β-lactamase-producing Enterobacteriaceae. Antimicrob Agents Chemother 45, 3548–3554 (2001).

27. H. R. Meredith, J. K. Srimani, A. J. Lee, A. J. Lopatkin, L. You, Collective antibiotic tolerance: mechanisms, dynamics and intervention. Nature Chemical Biology 11, 182–188 (2015).

28. L. Geyrhofer et al., Minimal Surviving Inoculum in Collective Antibiotic Resistance. mBio 14, e0245622 (2023).

29. T. Artemova, Y. Gerardin, C. Dudley, N. M. Vega, J. Gore, Isolated cell behavior drives the evolution of antibiotic resistance. Molecular systems biology 11, 822 (2015).

30. S. H. Lee et al., Association between type A blaZ gene polymorphism and cefazolin inoculum effect in methicillin-susceptible Staphylococcus aureus. Antimicrobial Agents and Chemotherapy 60, 6928–6932 (2016).

31. S. O. Lee, S. Lee, S. Park, J. E. Lee, S. H. Lee, The cefazolin inoculum effect and the presence of type A blaZ gene according to agr genotype in methicillin-susceptible Staphylococcus aureus bacteremia. Infection & Chemotherapy 51, 376 (2019).

32. E. C. Nannini et al., Determination of an inoculum effect with various cephalosporins among clinical isolates of methicillin-susceptible Staphylococcus aureus. Antimicrobial agents and chemotherapy 54, 2206–2208 (2010).

33. W. A. Craig, S. M. Bhavnani, P. G. Ambrose, The inoculum effect: fact or artifact? Diagnostic microbiology and infectious disease 50, 229–230 (2004).

34. D. Yong et al., Characterization of a new metallo-β-lactamase gene, bla NDM-1, and a novel erythromycin esterase gene carried on a unique genetic structure in Klebsiella pneumoniae sequence type 14 from India. Antimicrobial agents and chemotherapy 53, 5046–5054 (2009).

35. R. Lutz, H. Bujard, Independent and tight regulation of transcriptional units in Escherichia coli via the LacR/O, the TetR/O and AraC/I1-I2 regulatory elements. Nucleic acids research 25, 1203–1210 (1997).

36. A. U. Khan, L. Maryam, R. Zarrilli, Structure, genetics and worldwide spread of New Delhi metallo-β-lactamase (NDM): a threat to public health. BMC microbiology 17, 101 (2017).

37. W. P. Hempfling, S. E. Mainzer, Effects of varying the carbon source limiting growth on yield and maintenance characteristics of Escherichia coli in continuous culture. Journal of bacteriology 123, 1076–1087 (1975).

38. A. G. Marr, Growth rate of Escherichia coli. Microbiological reviews 55, 316–333 (1991).

39. O. Fridman, A. Goldberg, I. Ronin, N. Shoresh, N. Q. Balaban, Optimization of lag time underlies antibiotic tolerance in evolved bacterial populations. Nature 513, 418–421 (2014).

40. S. K. Gill et al., Increased airway glucose increases airway bacterial load in hyperglycaemia. Scientific reports 6, 27636 (2016).

41. J. Hosmer, A. G. McEwan, U. Kappler, Bacterial acetate metabolism and its influence on human epithelia. Emerging Topics in Life Sciences 8, 1–13 (2023).

42. H. Rytter et al., The pentose phosphate pathway constitutes a major metabolic hub in pathogenic Francisella. PLoS Pathogens 17, e1009326 (2021).

43. A. Grove, The delicate balance of bacterial purine homeostasis. Discover Bacteria 2, 14 (2025).

44. CLSI, Methods for dilution antimicrobial susceptibility tests for bacteria that grow aerobically; approved standard - tenth edition. . M07–A10 (2015).

45. S. Snoeck, C. Guidi, M. De Mey, “Metabolic burden” explained: stress symptoms and its related responses induced by (over) expression of (heterologous) proteins in Escherichia coli. Microbial Cell Factories 23, 96 (2024).

46. J. Weber, Z. Li, U. Rinas, Recombinant protein production provoked accumulation of ATP, fructose-1, 6-bisphosphate and pyruvate in E. coli K12 strain TG1. Microbial Cell Factories 20, 169 (2021).

47. J. H. Yang et al., Antibiotic-induced changes to the host metabolic environment inhibit drug efficacy and alter immune function. Cell host & microbe 22, 757–765. e753 (2017).

48. S. Meylan et al., Carbon sources tune antibiotic susceptibility in Pseudomonas aeruginosa via tricarboxylic acid cycle control. Cell chemical biology 24, 195–206 (2017).

49. A. Bourreau, V. Le Mabecque, A. Broquet, J. Caillon, Prevalence of a cefazolin inoculum effect associated with blaZ gene types, and clinical outcomes among methicillin-susceptible Staphylococcus aureus blood isolates of patients with infective endocarditis. Infectious Diseases Now 53, 104626 (2023).

50. B. Jean et al., β-Lactam Inoculum Effect in Methicillin-Susceptible Staphylococcus aureus Infective Endocarditis. JAMA Network Open 7, e2451353–e2451353 (2024).

51. W. R. Miller et al. (2018) The cefazolin inoculum effect is associated with increased mortality in methicillin-susceptible Staphylococcus aureus bacteremia. in Open forum infectious diseases (Oxford University Press US), p ofy123.

52. J. Guo et al., Implications of pH and Ionic Environment in Chronic Diabetic Wounds: An Overlooked Perspective. Clin Cosmet Investig Dermatol 17, 2669–2686 (2024).

53. R. Mann, D. G. Mediati, I. G. Duggin, E. J. Harry, A. L. Bottomley, Metabolic Adaptations of Uropathogenic E. coli in the Urinary Tract. Front Cell Infect Microbiol 7, 241 (2017).

54. A. Palomino et al., Metabolic genes on conjugative plasmids are highly prevalent in Escherichia coli and can protect against antibiotic treatment. The ISME Journal 17, 151–162 (2023).

55. R. J. Wallace, W. H. Holms, Maintenance coefficients and rates of turnover of cell material in Escherichia coli ML308 at different growth temperatures. FEMS Microbiol Lett 37, 317–320 (1986).

