## Supplementary Information for "A biphasic metabolic–growth trade-off governs β-lactam inoculum effects"

**Supplementary Figures and Figure Legends**

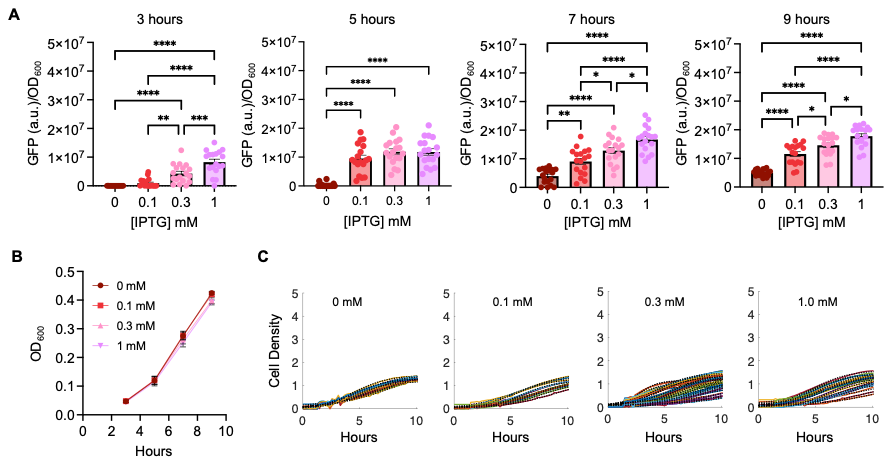

**Supplementary Figure 1: Raw data for Figure 1.**

A) Green fluorescent protein (GFP, arbitrary units, a.u.) normalized by cell density (OD_600_) as a function of [IPTG] and at different time points as noted above each graph. Error bars = SEM from 6 biological replicates per % CAA. ANOVA, P < 0.0001, Tukey HSD (** P = 0.0023, *** P = 0.0002, **** P <0.0001)
B) Cell density (OD) of cell cultures where GFP (panel A) was measured. From hours 5-9, growth was in log phase. Error bars = SEM from 6 biological reps per % CAA.
C) Raw growth curves (colored lines) and computed fits (black dotted lines) were used to determine the maximum growth rate. [IPTG] is indicated on each panel.

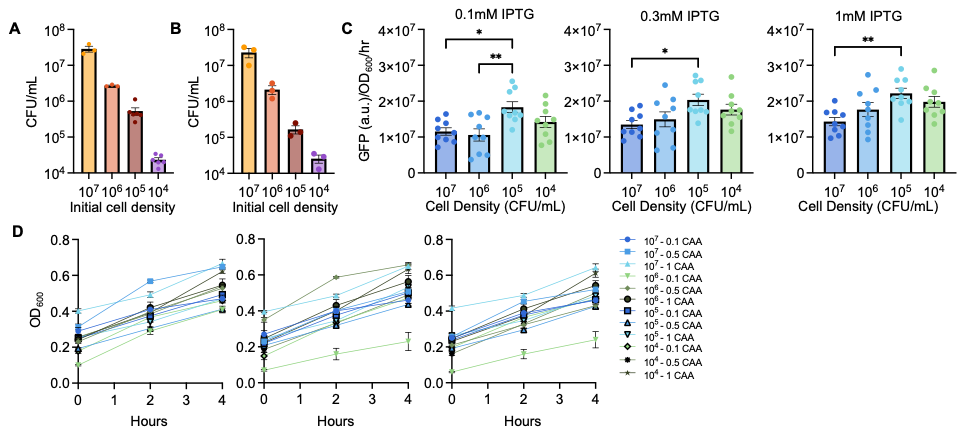

**Supplementary Figure 2: The rate of GFP expression as a function of initial cell density.**

A) Experimentally determined initial cell density of NDM-1 expressing *E. coli*. SEM from > 3 biological replicates.
B) Experimentally determined initial cell density of GFP-expressing *E. coli*. SEM from 3 biological replicates.
C) Rate of green fluorescent protein (GFP, arbitrary units, a.u.) expression normalized by cell density (OD_600_) per hour as a function of [IPTG] at different initial densities. The rate was determined by fitting a linear line through a plot of GFP/OD_600_ over 4 hours. Error bars = SEM from 3 biological replicates per % CAA. 0.1 mM: ANOVA, P = 0.0036, Tukey HSD, P *=0.0136, **=0.0041. 0.3 mM: ANOVA, P = 0.0255, Tukey HSD, * P = 0.0248. 1 mM: ANOVA, P = 0.0077, Tukey HSD, ** P = 0.0053.
D) Cell density (OD_600_) as a function of time for each initial cell density at all three % CAA. These plots demonstrate the trend in cell growth during which the GFP expression rate was determined. SEM from 3 biological replicates.

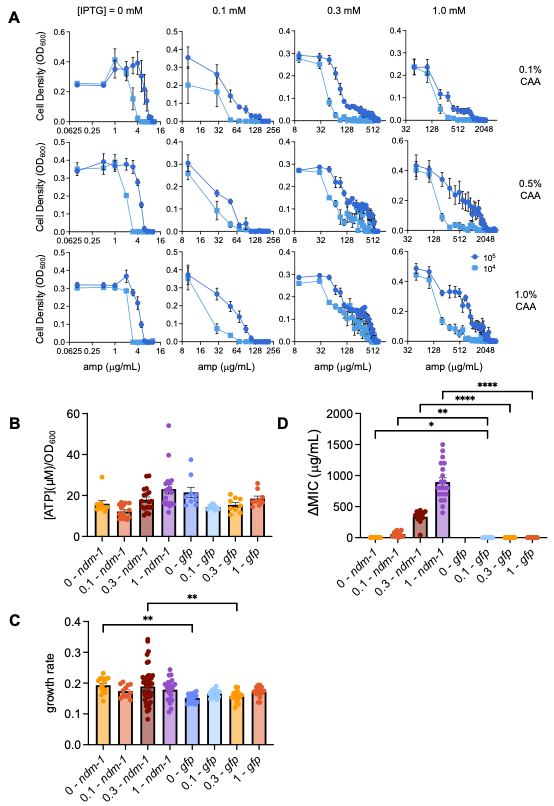

**Supplementary Figure 3: Raw MIC data for *E. coli* grown in media with glucose.**

A) Cell density (OD_600_) at 24 hours of 10^5^ (dark blue) and 10^4^ (light blue) CFU/mL initial density populations as a function of ampicillin (amp). [IPTG] as indicated. SEM from ≥ 5 biological replicates. SEM > 3 biological replicates for each % of casamino acids (CAA).
B) [ATP] per cell (OD_600_) of *E. coli* expressing either NDM-1 or GFP in medium with different [IPTG]. ANOVA, P = 0.0001, Sidak’s multiple comparison for each [IPTG], P ≥ 0.22. SEM from ≥ 3 biological replicates per %CAA.
C) Growth rate of *E. coli* expressing either NDM-1 or GFP in medium with different [IPTG]. ANOVA, P = 0.0027, Sidak’s multiple comparison for each [IPTG], P ** ≤ 0.0088. SEM from ≥ 3 biological replicates per %CAA
D) ΔMIC of *E. coli* expressing either NDM-1 or GFP in medium with different [IPTG]. Kruskal- Wallis, P < 0.0001, Dunn’s multiple comparison for each [IPTG] (* P = 0.034, ** P = 0.0033, **** P < 0.0001). SEM from ≥ 3 biological replicates per %CAA.

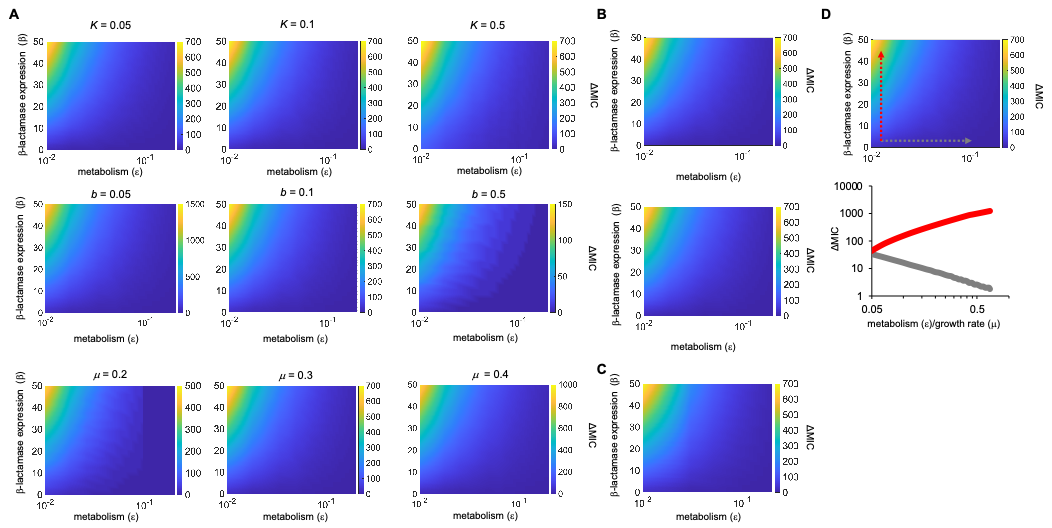

**Supplementary Figure 4: Sensitivity analysis of key parameters in our mathematical model (Eq. 1)**

A) Sensitivity of ΔMIC to perturbations in model parameters *β*, *μ*, *ε,* *K*, and *b*. Simulation time (*t*) = 24 hours. Parameter values consistent with Supplementary Table 2 unless otherwise noted in panel A.
B) Qualitative behavior of our modeling predictions persists at extended simulation times (top: t = 24 hours; bottom: t = 50 hours).
C) Removal of *ε* from $\left( \frac{N*A*\epsilon*b}{K+N*\beta} \right)$ does not impact the qualitative predictions of our model. *t* = 24 hours. Parameter values consistent with Supplementary Table 2.
D) The impact of individually increasing β-lactamase expression (β, red) or metabolism (ε, gray) on ΔMIC. Arrows illustrate general areas where slices of the heatmap (top) were used for simulations (bottom).

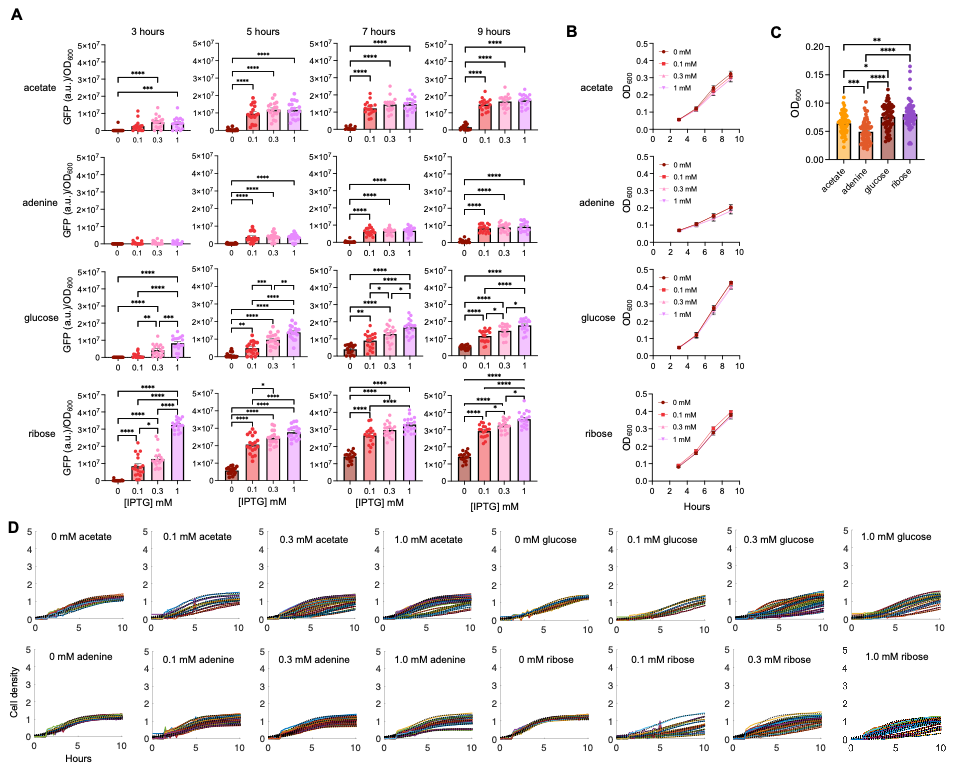

**Supplementary Figure 5: Raw data for Figure 3.**

A) Green fluorescent protein (GFP, arbitrary units, a.u.) normalized by cell density (OD_600_) as a function of [IPTG] and time as indicated in each column. SEM from 6 biological replicates per % of casamino acids. Carbon source as indicated. For all plots, ANOVA except for adenine – 3 hours, P < 0.0001. Tukey HSD (* P ≤ 0.0472, ** P = 0.0023 , *** P ≤ 0.0005, **** P < 0.0001 )
B) Cell density (OD) of cell cultures where GFP (panel A) was measured. From hours 5-9, growth was in log phase. Error bars = SEM from 6 biological reps per % CAA.
C) Average OD_600_ at which [ATP] was measured. Data averaged across all [IPTG] and % CAA for each carbon source. Kruskal Wallis, P < 0.0001, Dunn’s multiple comparison test (* P = 0.0379, ** P = 0.0014, *** P = 0.0009, **** P < 0.0001).
D) Raw growth curves were used to determine growth. Panels contain plots for all measured % CAA. Colored lines = experimental data. Black, dotted lines = modeling fit.

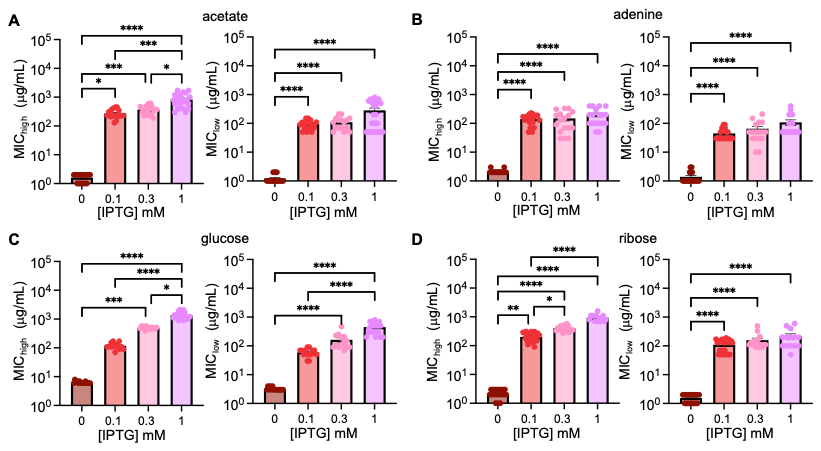

**Supplementary Figure 6: MIC of 10^5^ and 10^4^ CFU/mL initial density populations.** MIC of 10^5^ (left) and 10^4^ (right) CFU/mL initial density populations grown in acetate (A), adenine (B), glucose (C), and ribose (D). For all panels, Kruskal-Wallis, P < 0.0001. Dunn’s multiple comparison test (* P ≤ 0.0456, ** P = 0.0059, *** P ≤ 0.0002, **** P < 0.0001)

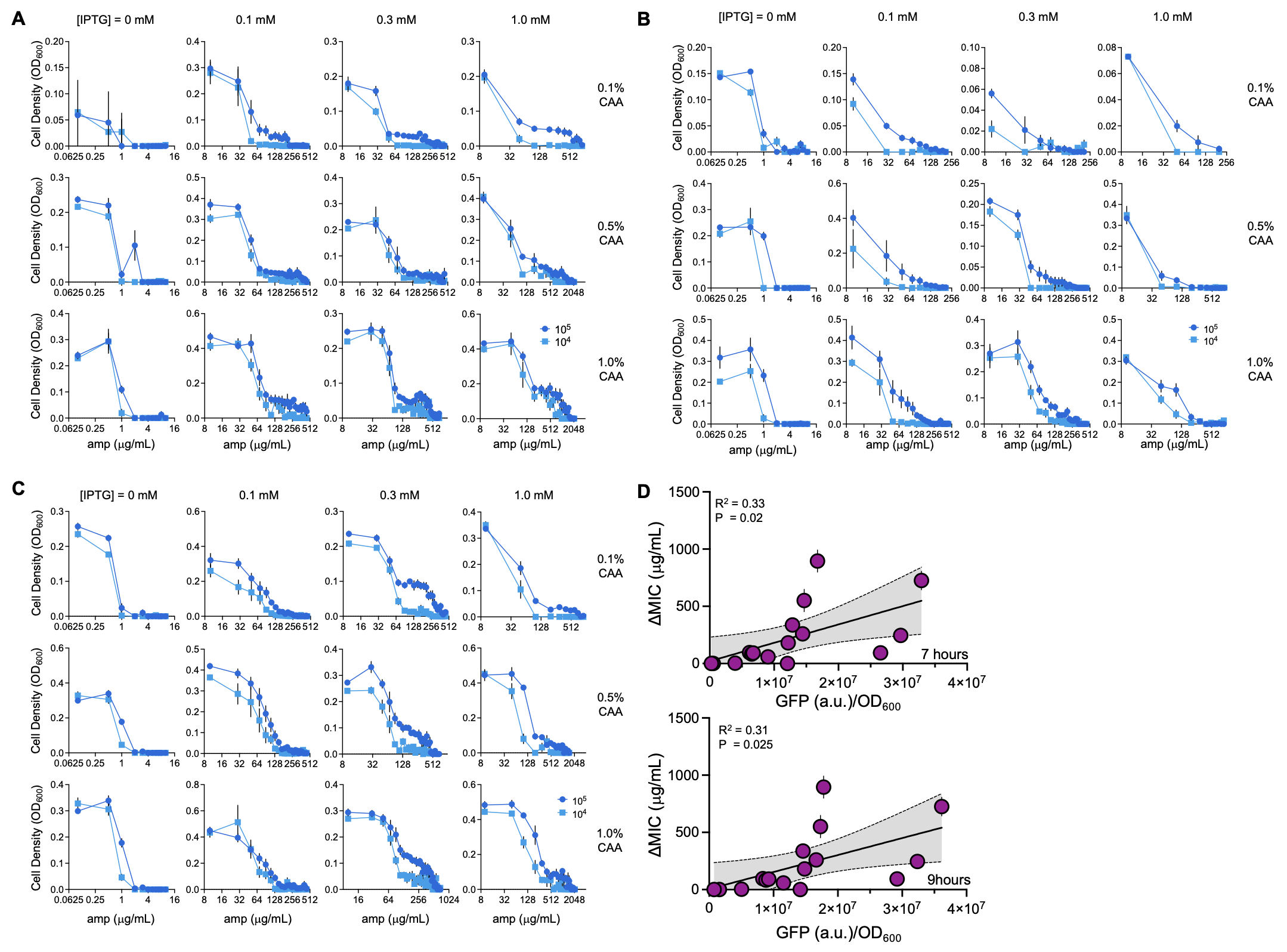

**Supplementary Figure 7: Raw data for Figure 4.**

A) Cell density at 24 hours of 10^5^ (dark blue) and 10^4^ (light blue) CFU/mL initial density populations at increasing [ampicillin] and grown in acetate as the carbon source. For panels A-C: SEM from ≥ 3 biological replicates. % of casamino acids (CAA) and [IPTG] indicated on the plot.
B) Cell density vs. ampicillin when adenine is used as the carbon source
C) Cell density vs. ampicillin when adenine is used as the carbon source
D) Regression analysis between ΔMIC of ampicillin and GFP/OD_600_ for all conditions. Time of GFP measurement indicated on the plot. R^2^ and P from simple linear regression. Grey = 95% confidence interval.

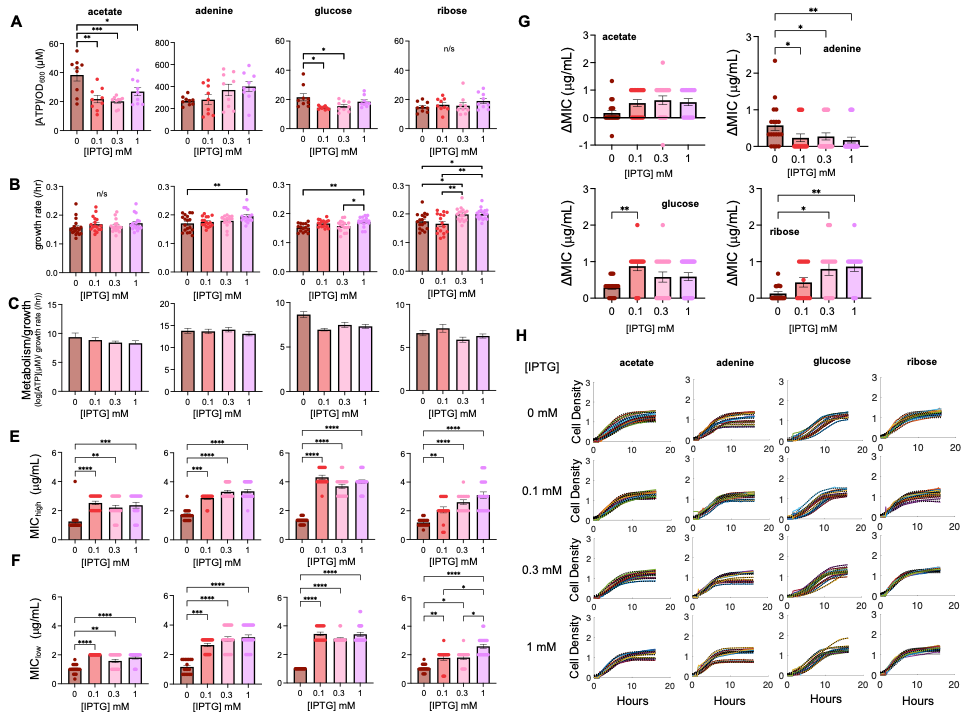

**Supplementary Figure 8: Physiological measures and ΔMIC of *E. coli* expressing GFP.**

A) [ATP] normalized by cell density (OD_600_) for GFP-expressing *E. coli* grown with different carbon sources and [IPTG]. SEM from 3 biological replicates per % of casamino acids (CAA).
B) Growth rate for GFP-expressing *E. coli* grown with different carbon sources and [IPTG].SEM from ≥ 5 replicates per % of casamino acids.
C) Metabolism normalized by growth rate for each combination of carbon source and [IPTG]. [ATP] and growth rate from panels A and B, respectively. Error bars = SEM.
D) Raw growth curves were used to determine growth. Panels contain plots for all % CAA measured. Colored lines = experimental data. Black, dotted lines = modeling fit.
E) MIC of 10^5^ CFU/mL initial density populations of *GFP*-expressing *E. coli.* SEM from > 5 biological replicates per % CAA. For all panels, Kruskal-Wallis P < 0.0001 (Dunn test ** P ≤ 0.0091, *** P ≤ 0.0005 , ****. P < 0.0001).
F) MIC of 10^4^ CFU/mL initial density populations of *GFP*-expressing *E. coli.* SEM from > 5 biological replicates per % CAA. For all panels, Kruskal-Wallis P < 0.0001 (Dunn test * P ≤ 0.0390, ** P ≤ 0.0093, *** P = 0.0001 , ****. P < 0.0001).
G) ΔMIC of *gfp*-expressing bacteria. ΔMIC calculated from data in panels E and F. Error bars = SEM.
H) Raw growth curves (colored lines) and computed fits (black dotted lines) were used to determine the maximum growth rate. [IPTG] is indicated on each panel.

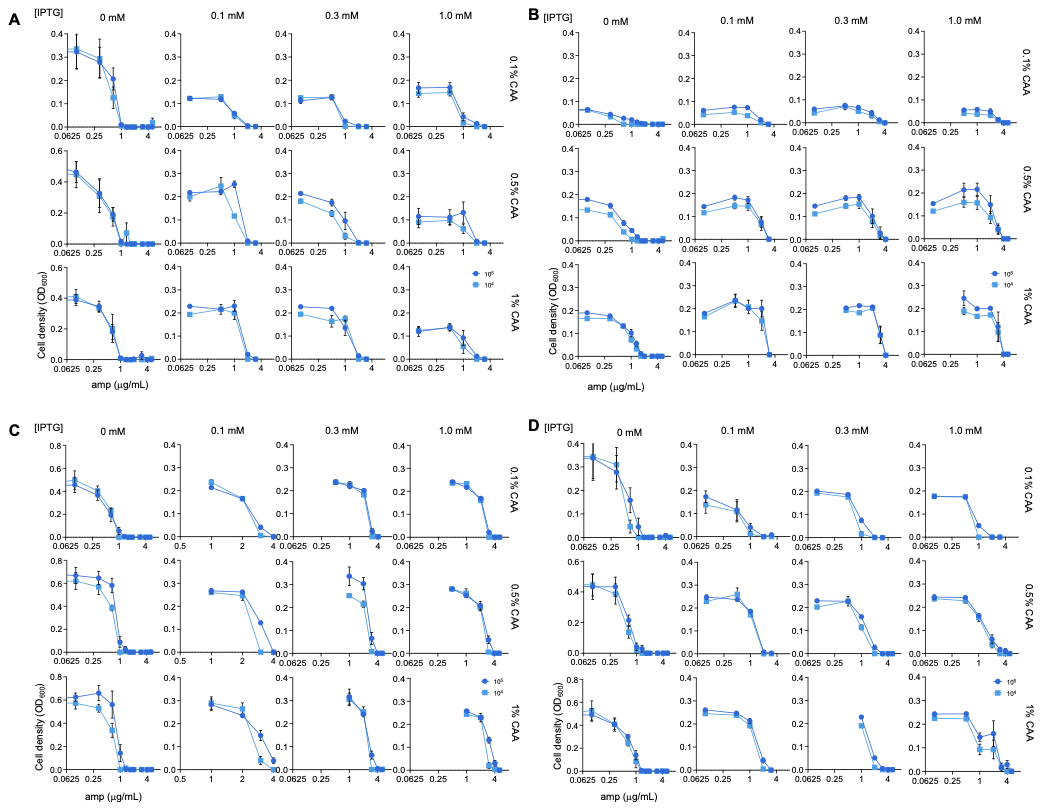

**Supplementary Figure 9: Raw data used to determine the MIC of *E. coli* expressing GFP.** Cell density at 24 hours of GFP expressing *E. coli* at 24 hours of 10^5^ (dark blue) and 10^5^ (light blue) initial density populations as a function of ampicillin (amp) grown in medium with acetate (A), adenine (B), glucose (C), and ribose (D). SEM from 6 biological replicates. [IPTG] and % of casamino acids (CAA) indicated on the plot.

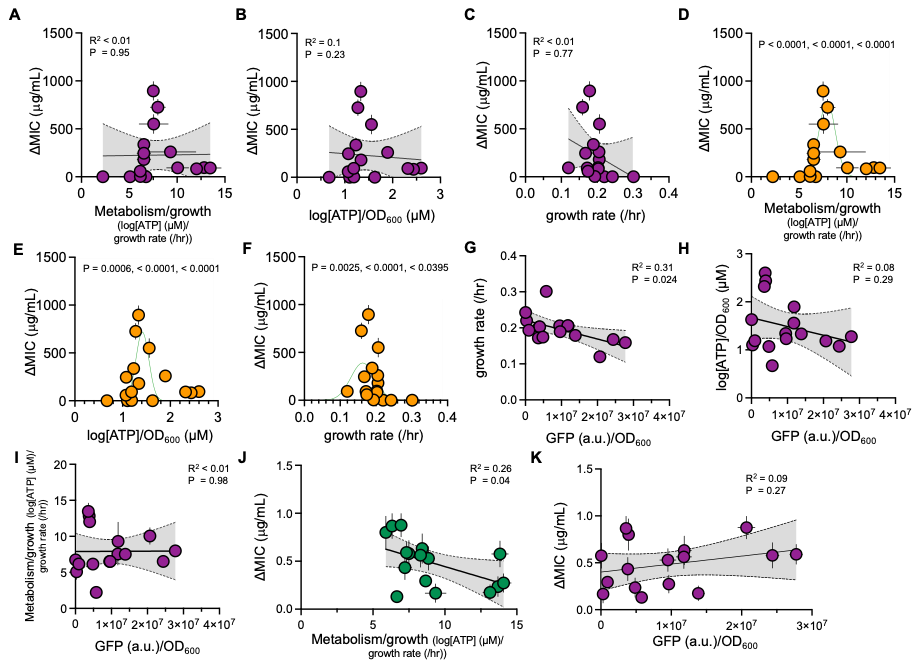

**Supplementary Figure 10: Linear regression analysis and Gaussian curve fits associated with Fig. 4.**

A) ΔMIC as a function of metabolism normalized by growth. For panels A-C, line, P values, and R^2^ value from a linear regression. ΔMIC from Fig. 4A; metabolism normalized by growth from Fig. 3F.
B) ΔMIC as a function of log[ATP]/OD_600_. [ATP] from Fig. 3D.
C) ΔMIC as a function of growth rate. Growth rate from Fig. 3E.
D) Gaussian curve fit through ΔMIC as a function of log[ATP]/OD_600_. For panels D-F, line, P values (in order – peak value, critical point, growth rate), and R^2^ value from a Gaussian fit.
E) Gaussian curve fit through ΔMIC as a function of growth rate—data from Fig. 3E.
F) Gaussian curve fit through ΔMIC as a function of GFP/OD_600_—data from Fig. 3C.
G) GFP/OD_600_ at 5 hours as a function of growth rate. For panels G-I, SEM from ≥ 4 biological replicates. R^2^ and P values from linear regression. GFP/OD_600_ from Fig. 3C; growth rate from Fig. 3E.
H) GFP/OD_600_ at 5 hours as a function of log[ATP]/OD_600_. log[ATP]/OD_600_ from Fig. 3D.
I) GFP/OD_600_ at 5 hours as a function of metabolism normalized by growth. metabolism normalized by growth from Fig. 3F.
J) ΔMIC as a function of metabolism normalized by growth for *E. coli* expressing NDM-1. Data includes 0 mM IPTG control conditions. ΔMIC from Fig. 4A; metabolism normalized by growth from Fig. 3F.
K) Linear regression of ΔMIC as a function of GFP/OD_600_ for *E. coli* expressing GFP. Data includes 0 mM IPTG control conditions.

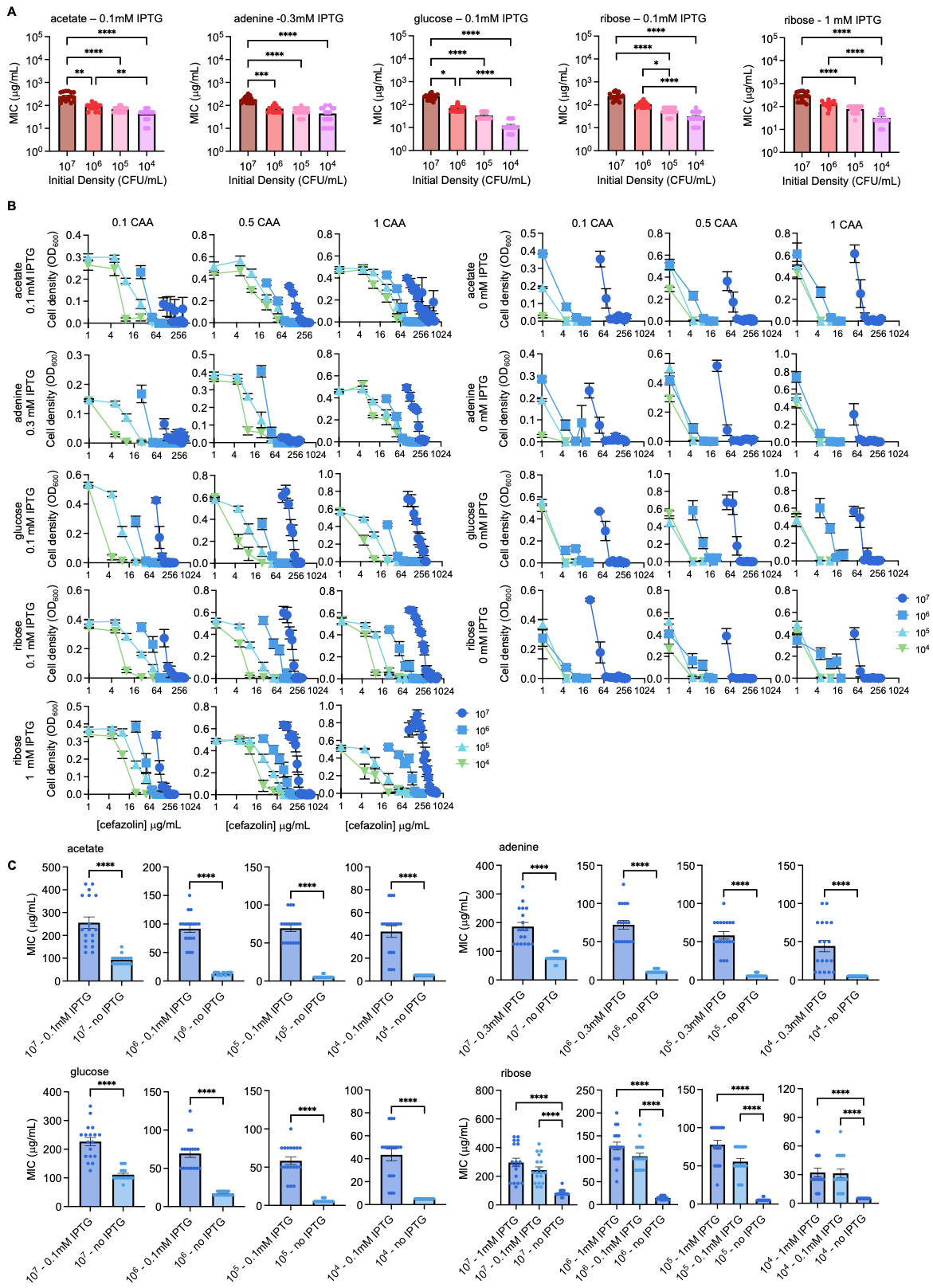

**Supplementary Figure 11: Raw MIC for *E. coli* grown in cefazolin.**

A) MIC as a function of initial density and using the carbon source and [IPTG] as indicated on each panel. For all panels, SEM from 6 biological replicates per % of casamino acids (CAA). For all panels, Kruskal-Wallis, P < 0.0001, Dunn’s multiple comparison test (* P ≤ 0.031, ** P ≤ 0.0034, *** P = 0.0002, **** P < 0.0001 )
B) Cell density at 24 hours of multiple initial density populations (10^7^, 10^6^, 10^5^, 10^4^ CFU/mL) at increasing [cefazolin]. Carbon source and % CAA are indicated on the plot. For panels A-C: SEM from ≥ 3 biological replicates. % of casamino acids (CAA) and [IPTG] indicated on the plot.
C) MIC of induced and uninduced (0 mM IPTG) cultures grown in the [IPTG], carbon source, and initial density as indicated on the plot. For all panels, Mann-Whitney test, **** P < 0.0001.

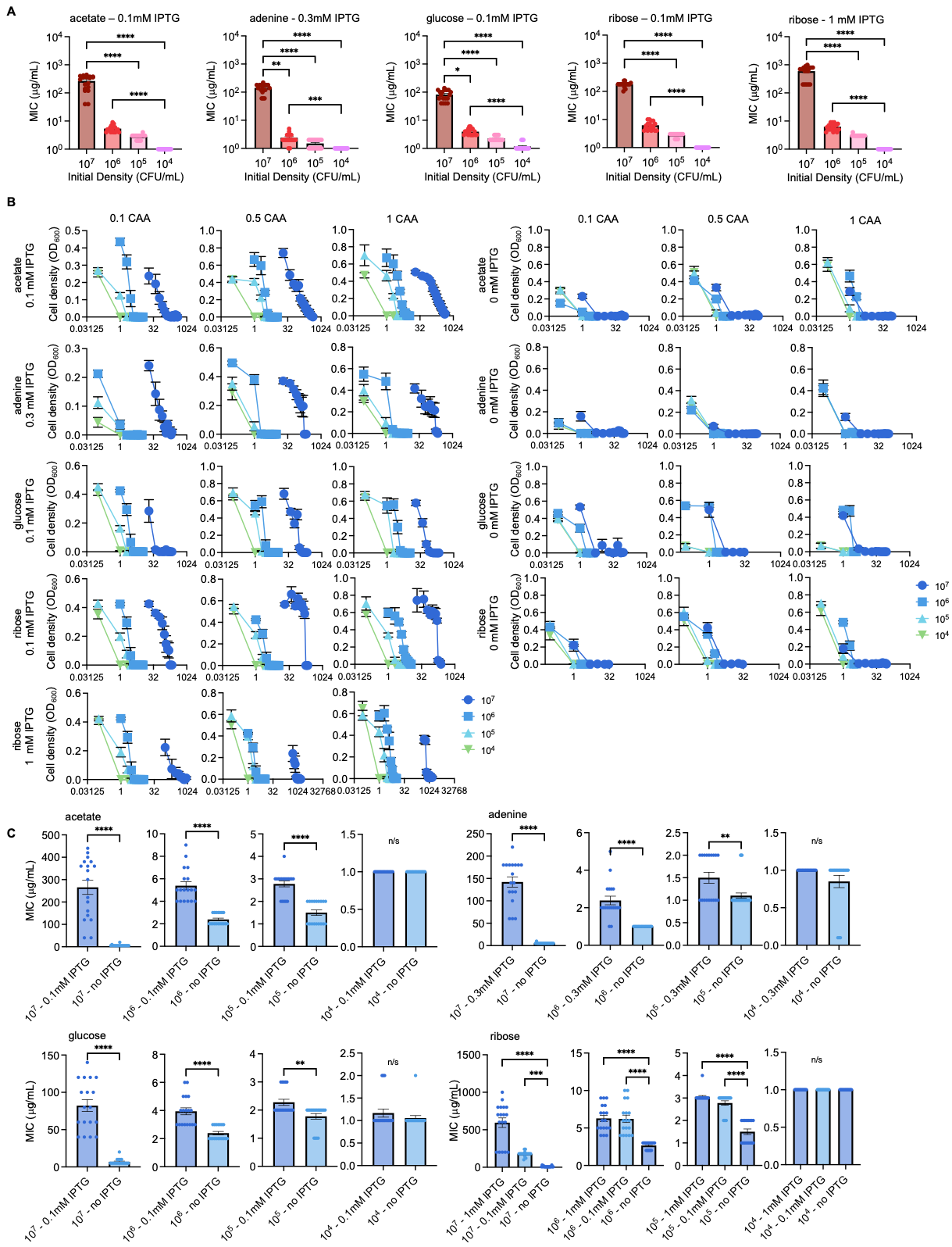

**Supplementary Figure 12: Raw MIC for *E. coli* grown in imipenem.**

A) MIC as a function of initial density and using the carbon source and [IPTG] as indicated on each panel. For all panels, SEM from 6 biological replicates per % of casamino acids (CAA). For all panels, Kruskal-Wallis, P < 0.0001, Dunn’s multiple comparison test (* P = 0.0329, ** P = 0.0034, *** P = 0.0008, **** P < 0.0001 )
B) Cell density at 24 hours of multiple initial density populations (10^7^, 10^6^, 10^5^, 10^4^ CFU/mL) at increasing [imipenem]. Carbon source and % CAA are indicated on the plot. For panels A-C: SEM from ≥ 3 biological replicates. % of casamino acids (CAA) and [IPTG] indicated on the plot.
C) MIC of induced and uninduced (0 mM IPTG) cultures grown in the [IPTG], carbon source, and initial density as indicated on the plot. For all panels, Mann-Whitney test, ** P ≤ 0.0048. *** P = 0.0006, **** P < 0.0001.

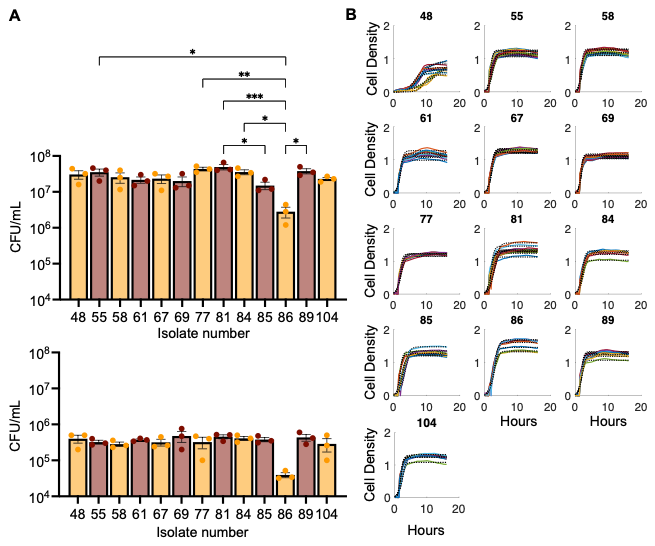

**Supplementary Figure 13: Physiological measures of β-lactamase-expressing E. coli clinical isolates.**

A) Experimentally determined initial densities (10^7^ top, 10^5^ bottom) of *E. coli* clinical isolates. SEM from 3 biological replicates. ANOVA P = 0.0009, *** P = 0.0006, ** P = 0.0035, * P ≤ 0.0305, Tukey HSD.
B) Raw growth curves were used to determine the growth rates of clinical isolates, as indicated in the plot. Colored lines = experimental data. Black, dotted lines = modeling fit.

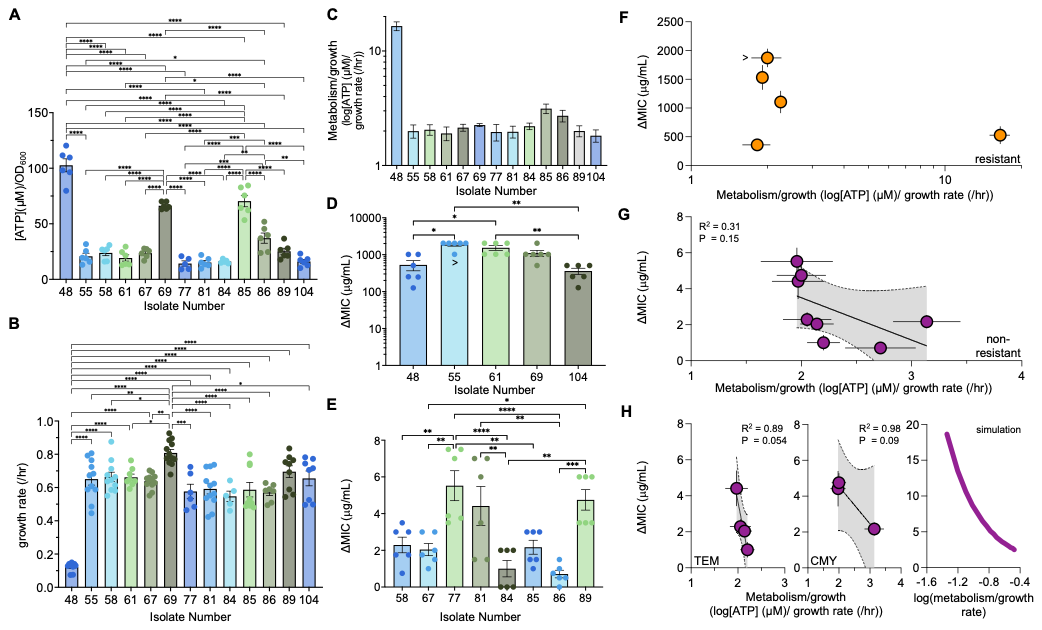

**Supplementary Figure 14: Raw MIC for *E. coli* clinical isolates.**

A) MIC of imipenem for high-density populations (~10^7^ CFU/mL) of *E. coli* clinical isolates. SEM from 6 biological replicates. Kruskal-Wallis, P < 0.0001, Dunn’s multiple comparison test ( * P ≤ 0.035, ** P ≤ 0.0091, *** P ≤ 0.0007, **** P < 0.0001). For isolate 55, (panels D and E) > indicates that ΔMIC is greater than the value reported, as some replicates showed growth at their highest concentration of imipenem that could be tested
B) MIC of imipenem for low-density populations (~10^5^ CFU/mL) of *E. coli* clinical isolates. SEM from 6 biological replicates. Kruskal-Wallis, P < 0.0001, Dunn’s multiple comparison test ( * P ≤ 0.0191, ** P ≤ 0.0075).
C) Raw cell density after 24 hours of E. coli clinical isolates challenged with increasing concentrations of imipenem. SEM from 6 biological replicates.
D) Linear regression of ΔMIC of imipenem as a function of log(ATP)/growth rate for *E. coli* clinical strains expressing TEM-type β-lactamases without isolate 86.

**Supplementary Table 1: Average residual values for growth rates presented in Figure 1.** %CAA = casamino acids. SEM from ≥ 3 biological replicates.

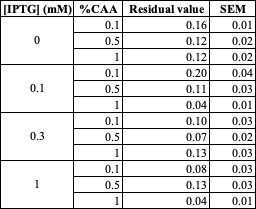

**Supplementary Table 2:** Parameters and their values used in our mathematical model (Eq. 1). For additional justification, see *Methods*.

| Parameter | Value | Justification |
| --- | --- | --- |
| Growth rate (*μ*) | 0.3/hr | Approximates highest experimental growth rate. |
| Metabolism (*ε*) | .01 mmol/g/hr | Smallest reported maintenance coefficients for *E. coli* |
| β-lactamase expression (*β*) | 1-300 (unitless) | Estimated based on fold induction of the P_lac-ara-1_ promoter. |
| Antibiotic-specific death rate (*b*) | 0.1/hr | Estimated from reported antibiotic survival data. |
| Half maximal killing rate of the antibiotic (*K*) | 0.1 | Estimated to fit experimental data |
| Cell density (*C)* | *C_high_* =0.05  *C_low_* = 0.0001 | Estimated using experimental initial densities. |
| Carrying capacity (*C_m_*) | 1 (unitless) | Normalized to cell density (*C*) |

**Supplementary Table 3: Average residual values for growth rates presented in Figure 3.** CAA = casamino acids. SEM from ≥ 3 biological replicates.

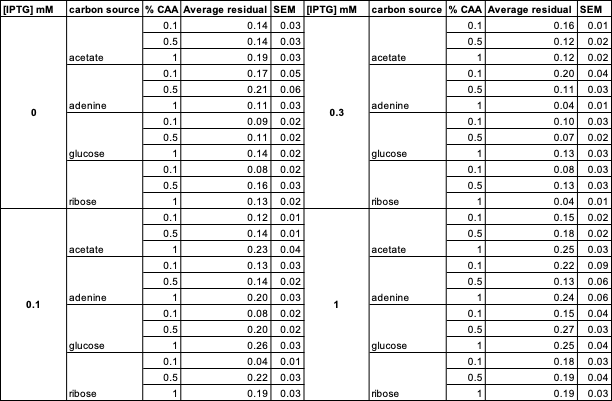

**Supplementary Table 4: Average residual values for growth rates of GFP-expressing *E. coli* presented in Supplementary Figure 8.** CAA = casamino acids. SEM from ≥4 biological replicates.

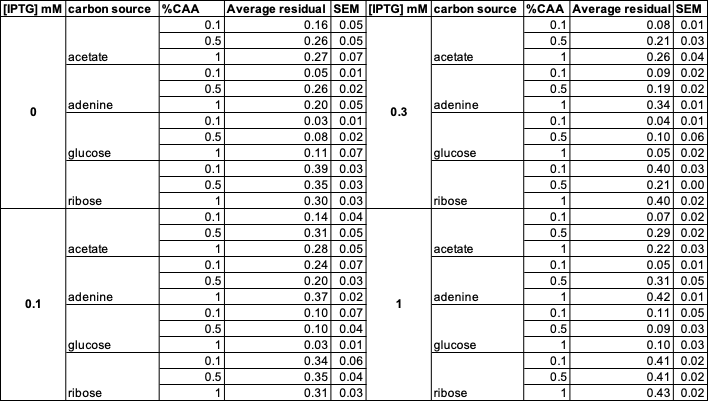

**Supplementary Table 5: Characteristics of *E. coli* clinical strains expressing different β-lactamases.** Susceptible/resistant status obtained from the Biosample Accession number listed for each strain.

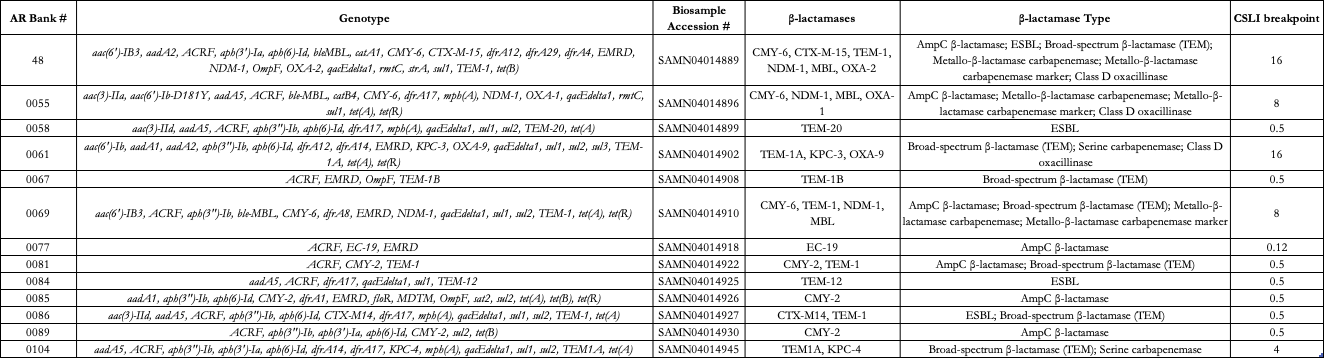

**Supplementary Table 6: Average residual values for growth rates of clinical strains of *E. coli* presented in Figure 6.** SEM from ≥ 5 biological replicates.

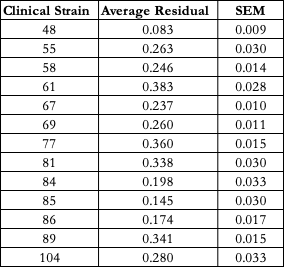
